# Astrocyte fatty acid uptake and ketogenesis in the mediobasal hypothalamus coordinate adaptive metabolic responses against high-fat diet exposure in male rodents

**DOI:** 10.64898/2026.10.07.757016

**Authors:** Keila Navarro i Batista, Nadine Ellersiek, Patricia Kulka, Gaia Serra, Francesco Prisco, Christelle Le Foll

## Abstract

The mediobasal hypothalamus (MBH) constitutes a central regulator of energy homeostasis; specialized neurons in this region integrate peripheral metabolic signals to coordinate processes like food intake, energy expenditure or glucose homeostasis. While MBH neuronal dysfunction has been linked to obesity, increasing evidence indicates that astrocytes in this region are also active participants in the modulation of energy balance. MBH astrocyte ketone body signaling, in particular, has been proposed as an adaptive mechanism aimed at regulating energy homeostasis during dietary challenge.

Here, we investigated the effects of MBH astrocytic lipid metabolism on the control of energy balance. We targeted distinct components of this pathway, ablating fatty acid (FA) transporter FATP4 in mice and knocking down 3-hydroxy-3-methylglutaryl-CoA lyase (HMGCL) in rats. MBH astrocyte FATP4 depletion markedly exacerbated high-fat diet (HFD)-induced obesity, increasing body weight and fat mass independently of food intake. Rather, this phenotype was driven by reduced energy expenditure and impaired brown adipose tissue oxidation. HMGCL knockdown in MBH astrocytes also promoted HFD-induced weight gain, not only through reduced energy expenditure but also through a transient hyperphagic response during early HFD exposure. *In vitro* studies in primary astrocyte cultures confirmed that disruption of FATP4 altered the expression of genes involved in mitochondrial lipid oxidation and ketogenesis. Similarly, HMGCL knockdown reduced the expression of monocarboxylate transporter MCT4, suggesting impaired astrocytic lactate and/or ketone body release.

Together, these findings identify MBH astrocyte lipid metabolism as a critical component of the hypothalamic circuitry regulating energy balance, highlighting astrocytes as active contributors to the pathogenesis of obesity.

**Graphical abstract:** 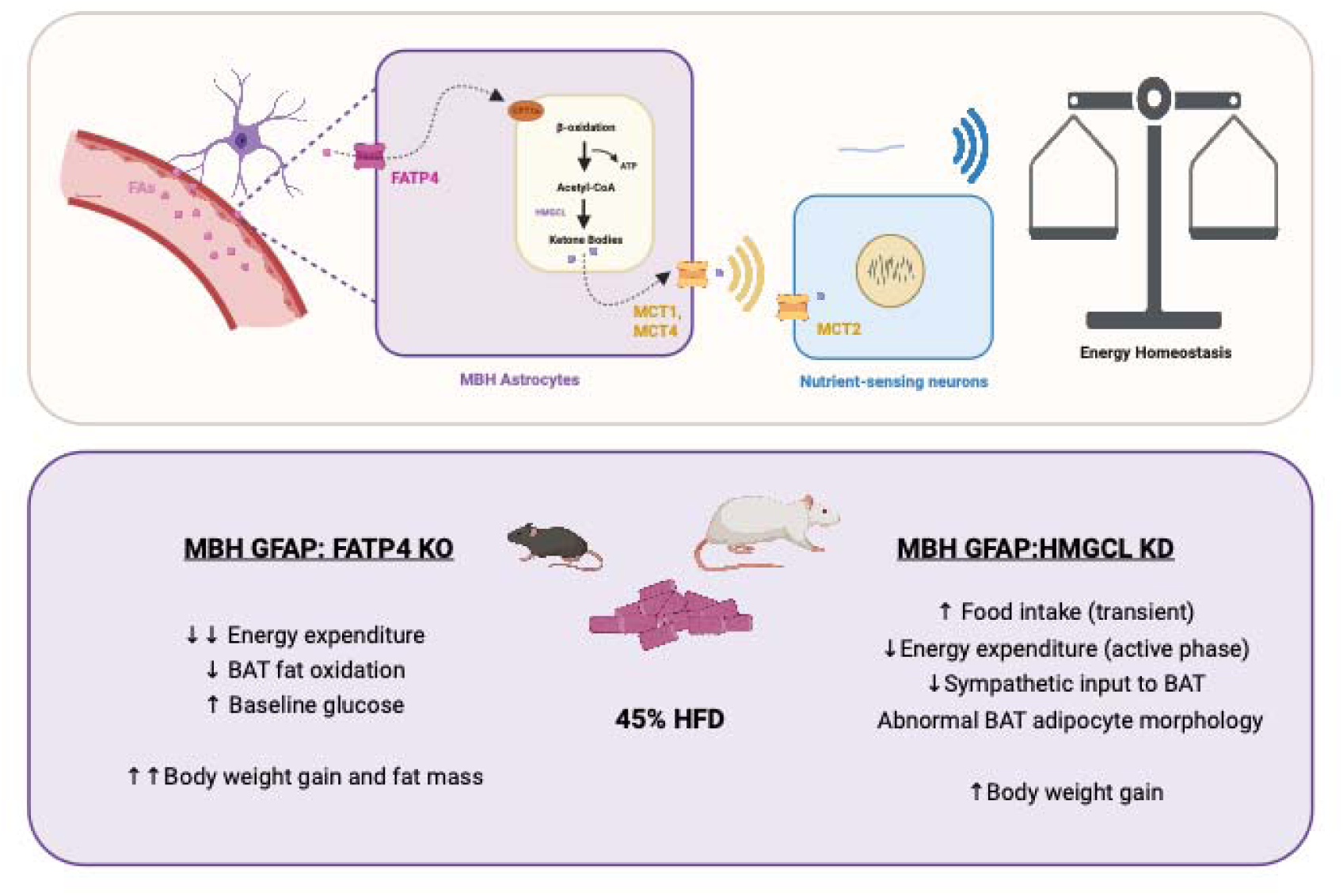

**Highlights:** • MBH astrocyte FATP4 depletion exacerbates HFD-induced obesity in mice

- Astrocyte HMGCL knockdown impairs metabolic adaptation to HFD in rats
- Astrocyte lipid metabolism supports energy expenditure and hypothalamic ketone signaling

## 1. Introduction

The mediobasal hypothalamus (MBH) serves as a central regulator of whole-body energy homeostasis [1]. Specialized neurons within this region sense and integrate peripheral metabolic cues to coordinate vital physiological processes such as food intake, energy expenditure or systemic glucose homeostasis [1–3]. Defects at the level of the MBH are widely recognized as major contributors to the onset and progression of numerous metabolic disorders, including obesity [4]. In both obese patients and animal models of the disease, MBH dysfunction is typically associated with altered activity of nutrient-sensing neurons such as pro-opiomelanocortin (POMC) and agouti-related peptide (AgRP) neurons [5–8]. However, it also involves pathological remodeling of non-neuronal populations, including astrocytes [9].

Astrocytes are key modulators of neuronal activity within the MBH by regulating synaptic transmission and nutrient sensing, ultimately supporting the control of energy balance [9,10]. These functions may be of relevance under conditions of metabolic stress. The MBH is highly sensitive to increased fatty acid (FA) availability, and while neuronal responses to FA have been well studied [11,12], accumulating evidence indicates that astrocyte activation also plays a critical role in maintaining homeostasis during dietary challenge [13,14]. Notably, whereas chronic astrogliosis has been associated with the pathogenesis of obesity, acute astrogliosis within the MBH in response to HFD exposure has been proposed to exert protective effects [13,14]. Consistent with this notion, disruption of acute astrogliosis exacerbated the effects of high-fat diet (HFD) exposure, leading to increased food intake [13].

Emerging evidence suggests that astrocyte lipid metabolism may also contribute to metabolic adaptation to HFD, particularly through ketone body production [15–17]. Several studies have reported that ketone body signaling can modulate the central control of energy homeostasis [18,19] [15–17]. However the reported effects of ketone bodies on food intake and metabolism remain inconsistent, so their physiological relevance is still under debate.

Ketone body infusion studies, administered either through the carotid or directly into the hypothalamus, have suggested that central ketone body signaling may promote a positive energy balance [18,19]. Specifically, 24-hour intracarotid infusion of ketone bodies has been reported to increase food intake in chow-fed mice, accompanied by elevated hypothalamic expression of the orexigenic neuropeptides NPY and AgRP [18]. Hypothalamic ketone body infusion over a 7-day period was shown to increase body weight gain, an effect attributed to reduced energy expenditure during the active phase [19]. Interestingly, hypothalamic ketone body delivery was also associated with downregulation of the monocarboxylate transporters MCT1 and MCT2, potentially reflecting a counter-regulatory response to sustained central hyperketonemia [19]. In this context, chow-fed mice heterozygous for MCT1 deficiency, resulting in partially impaired ketone body brain trafficking, were not resistant to ketone body infusion per se, but instead exhibited reduced susceptibility to weight gain [19].

In fasted rats, acute HFD consumption increased ketone bodies levels in the ventromedial hypothalamus (VMH) one of the nuclei of the MBH [17]. This response was found to precede HFD-induced reductions in food intake [17]. Interestingly, selective inhibition of ketone body production within the VMH had no effect on low-fat diet (LFD)-fed rats but significantly reversed anorectic responses to HFD, promoting hyperphagia [17]. Diet-resistant (DR) rats have been found to normalize their food intake three days after HFD exposure, whereas diet-induced obese (DIO) rats continued to overeat [15]. In DR rats, the reduction in food intake were once again disrupted by inhibition of ketogenesis [15]. Contrary to infusion experiments, these findings support a model in which endogenous ketone body within the MBH act as a negative feedback signal, limiting HFD-induced hyperphagia, rather than promoting an obesogenic phenotype. Prompted by these findings, we sought to determine how ketone body signaling may contribute to hypothalamic regulation of energy balance. Astrocytes represent the only source of ketogenesis in the central nervous system and increase ketone body synthesis in response to FA excess, making them likely drivers of HFD-induced ketogenesis in MBH [16,20]. We hypothesize that astrocyte ketogenesis and signaling, together with acute MBH astrogliosis, constitute an essential component of the adaptive response to HFD challenge.

Using complementary *in vivo* and *in vitro* approaches, we selectively manipulated astrocyte FA uptake and ketogenic capacity within the MBH. Specifically, we disrupted expression of the FA transporter protein 4 (FATP4) and the ketogenic enzyme 3-hydroxy-3-methylglutaryl CoA lyase (HMGCL) in MBH astrocytes to assess their roles in energy homeostasis under dietary stress aiming at establishing astrocyte lipid metabolism and ketone body signaling as central mediators of astrocyte-neuron communication in the MBH, with critical implications for the control of systemic energy homeostasis and metabolic health.

## 2. Materials and methods

### 2.1. MBH primary astrocytes studies

#### Primary astrocyte isolation

Primary astrocytes were isolated from 21-27-day-old C57BL/6 mice or Sprague-Dawley rats as previously described [21,22]. Briefly, MBH tissue was dissected, mechanically dissociated, and cultured on poly-D-lysine-coated flasks in supplemented Neurobasal-A medium. After 14 days *in vitro*, oligodendrocytes and microglia were removed by orbital shaking, and adherent astrocytes were detached with trypsin-EDTA and replated at 90,000 cells/well for subsequent experiments.

#### Astrocyte siRNA transfection and treatments

Astrocytes (approximately 70% confluence) were transfected with pooled siRNAs targeting Slc27a4 (FATP4) (Nr. L-063631-01-0005; Horizon Discovery Biosciences) or Hmgcl (Sequences derived from Hmgcl AAV Table S2; Microsynth) using Lipofectamine RNAiMAX (Thermo Fisher Scientific). Non-targeting siRNA served as a control (Nr. D-001210-01-05; Horizon Discovery Biosciences). Following transfection, cells were cultured in serum-free Neurobasal-A containing 2.5 mM glucose and treated with vehicle, palmitate or oleate (20 μM; Nr. 29556, 29557, 29558; Cayman Chemical) for up to 72 hours. Culture supernatants were collected every 6 or 12 hours for ketone body and lactate analysis, after which cells were harvested for RNA extraction. Detailed siRNA sequences are provided in the Supplementary Materials (Table S5).

To verify knockdown efficiency, Slc27a4 (FATP4) expression was quantified by qPCR as described in section 2.3. Extracellular L-lactate concentrations were determined using the Lactate Assay Kit II (Nr. MAK065; Sigma-Aldrich) according to the manufacturer’s instructions. Similarly, ketone body β-hydroxybutyrate concentrations were quantified (Autokit 3-HB; Fujifilm Wako Chemicals). Absorbance was measured at 450 nm and 405 nm respectively, using an Epoch 2 microplate reader (BioTek).

#### Visualization of FA uptake using the BODIPY-PA assay

Primary astrocytes were transfected with Slc27a4 (FATP4)-targeting or control siRNA as described above. After 48 hours, cells were incubated with the fluorescent palmitate (PA) analogue BODIPY-PA (10 μM; Nr. D3821, Thermo Fisher Scientific) for 5, or 30 minutes, washed and fixed with 4% paraformaldehyde, and immunostained for GFAP (1:500, Nr. ab53554, Abcam). Confocal images (Leica SP8 CLSM; Center for Microscopy and Image Analysis, University of Zurich) were acquired and processed in Fiji software (ImageJ 2.1.0). FA uptake was quantified by measuring the BODIPY signal colocalized with GFAP-positive astrocytes using Imaris software (Bitplane) in three separate wells.

### 2.2. MBH primary neuron isolation

Primary MBH neurons were isolated from 21-27-day-old C57BL/6 mice as previously described [23]. Briefly, hypothalamic slices containing the MBH were prepared, enzymatically dissociated with papain, mechanically triturated, and purified by BSA-gradient centrifugation. Freshly isolated neurons were plated for short-term culture before RNA extraction.

### 2.3. Gene expression analysis

Total RNA was extracted from astrocytes and freshly isolated neurons using the ReliaPrep RNA Cell Miniprep System (Nr. Z6011; Promega). cDNA was synthesized using the SensiFAST cDNA Synthesis Kit (Nr. BIO65054; Meridian Bioscience), and gene expression was quantified by qPCR using validated TaqMan assays.

Astrocyte gene expression was analyzed using the ΔΔCt method, whereas neuronal gene expression was quantified using standard curves generated from serial dilutions of pooled neuronal cDNA. Expression values were normalized to the mathematical mean of Cyclophilin or Gapdh and Rsp18. Primer and assay information is provided in the Supplementary Materials (Table S6).

### 2.4. Animal models and husbandry

All animal procedures were performed at the University of Zurich and approved by the Veterinary Office of the Canton of Zurich (license 247/2019, 071/2023 and 85/2023).

Male FATP4^flox/flox^ mice with a C57Bl/6 background [24] obtained from Walee Chamulitrat [25,26] were bred in-house and genotyped following standard protocols (primer sequences are provided in Table S1). Mice were group-housed under controlled conditions (22°C; 12h light-dark cycle (lights off at 11:00) with *ad libitum* access to standard chow (Diet 3336, Kliba Nafag) and water. Animals were 10-14 weeks old at the start of experiments and were allocated to experimental groups using weight-stratified randomization.

Male Sprague Dawley rats (Janvier Labs; 200-250g upon arrival) were acclimatized for one week under identical housing conditions and assigned to experimental groups using weight-stratified randomization. Animals were group-housed unless otherwise specified.

### 2.5. Weekly body weight and food intake monitoring

Body weight and food intake were measured twice weekly at the end of the light phase. Two weeks after stereotaxic surgery, animals were switched to a 45% high-fat diet (HFD; 4.73 kcal/g; macronutrient composition: 20% protein, 35% carbohydrates, 45% fat; E15744-344 SSNIFF Spezialdiäten GmbH,; FA composition described in [27]). Body weight at the time of HFD initiation was defined as baseline. Weekly body weight gain, caloric intake (kcal = food weight consumption per week x 4.73 kcal/g) and cumulative intake were subsequently calculated. Unless specified, mice and rats followed the same dietary protocol.

### 2.6. Stereotaxic viral delivery to the MBH

Astrocyte-specific gene manipulation in the MBH was achieved via bilateral stereotaxic injection of adeno-associated viral (AAV) vectors. Mice received AAV5/2 vectors encoding hGFAP-driven EGFP together with either Cre recombinase (FATP4 knockout) or EGFP alone (control), whereas rats received AAV9/2 vectors expressing hGFAP-driven EGFP together with either four shRNAs targeting Hmgcl or non-targeting shRNAs (control) (Table S2).

Animals underwent stereotaxic surgery under species-specific anesthesia and analgesia protocols (Table S2). Viral suspensions were injected bilaterally into the MBH at a rate of 100 nL/min using a 10 μL Hamilton syringe fitted with a 35G needle and controlled by a microsyringe pump (World Precision Instruments). Following injection, the needle remained in place for 10 minutes to minimize reflux before withdrawal. Animals received standard post-operative monitoring and analgesia for at least 72 hours.

### 2.7. Indirect calorimetry

Indirect calorimetry was performed using either the Promethion system (Sable Systems) or the Phenomaster system (TSE Systems), depending on the experimental paradigm and equipment availability. Energy expenditure (EE) and respiratory exchange ratio (RER) were calculated from O2 consumption and CO2 production using manufacturer software and standard equations [28]. Data were analyzed using CalR [29]. Hourly averaged data from 2-3 consecutive days were used to determine baseline EE and RER across 12 h-dark phase, 12 h-light phase, and the 24 h-total. To account for differences in body composition, EE was normalized to individual lean and fat mass using the formula: EE (kcal/h) / [lean body mass (g) + (0.2 × fat body mass (g)] [30]. In addition to EE and RER, the system continuously recorded food intake, allowing for hourly monitoring of feeding behavior.

#### Early HFD exposure in FATP knockout mice (Figure 3)

One week after stereotaxic surgery, a subset of mice was transferred to the Promethion indirect calorimetry system and allowed to acclimatize for one week under chow feeding. Once body weight and food intake had stabilized, animals were switched to a 45% HFD and monitored continuously until euthanasia.

#### Chronic HFD exposure in FATP knockout mice or HMGCL knockdown rats (Figures 4 and 7)

After five weeks of HFD, mice and rats were transferred to Phenomaster indirect calorimetry cages. Following acclimatization (5 days), EE and RER were recorded under *ad libitum* feeding. A subset of animals subsequently underwent a 12-hour fasting/refeeding protocol. In rats, an additional cohort was exposed sequentially to room temperature (22°C), thermoneutrality (30°C), and cold (10°C) to assess thermogenic responses. The first day of cold exposure served as habituation; therefore, only data from the last two days were included in the analysis.

### 2.8. Telemetry probe implantation

A subset of animals underwent surgical implantation of telemetry probes for continuous monitoring of locomotor activity and core body temperature. TA-F10 and TA-F40 (Data Sciences International) were implanted in mice and rats, respectively, four weeks after HFD initiation. Surgeries were performed under isoflurane anesthesia using the same peri-operative procedures described for stereotaxic surgery, except that the mice received long-acting buprenorphine for analgesia. Receiver plates connected to Ponemah software (Data Sciences International, Harvard Apparatus) were positioned beneath the indirect calorimetry cages, and telemetry data were acquitted continuously at 1-minute intervals.

### 2.9. Oral glucose tolerance test

To assess the impact of FATP4 knockout on peripheral glucose homeostasis, an oral glucose tolerance test (OGTT) was performed in mice eight weeks after HFD switch. Food was removed 2 h prior to lights off, and mice were gavaged (2 g/kg glucose) at lights off. Tail blood glucose was measured using a handheld glucometer (Contour XT; Ascensia Diabetes Care) before and 15, 30, 45, 60, 90, and 120 min after glucose administration. The area under the curve (AUC) from 0 to 120 minutes was calculated relative to each animal’s individual baseline (time 0).

### 2.10. Body composition analysis

Lean and fat mass were measured post-mortem using EchoMRI (EchoMRI Whole Body Composition Analyzer; Echo Medical Services) Fat mass (FM) and lean mass (LM) were calculated as the mean of two scans per animal and expressed in grams or as a percentage of total body weight.

### 2.11. Euthanasia

Animals were fasted for 2 hours and euthanized at the onset of the dark phase via intraperitoneal injection of pentobarbital (100-200 mg/kg, i.p.; Cantonal Pharmacy of Zurich, Switzerland). Blood, liver, and adipose tissue samples-including interscapular brown adipose tissue (iBAT), epididymal white adipose tissue (eWAT), and inguinal white adipose tissue (iWAT)-were collected. Brains intended for gene expression analyses were rapidly dissected and flash frozen. For immunohistochemistry, animals were either perfused after a 12-hour fast or 90 minutes following refeeding. Transcardial perfusion was performed with 0.1M phosphate buffer for 1 minute, followed by fixation with 4% paraformaldehyde (PFA) for 1.5 minutes. Brains were post-fixed in 4% PFA for 24 hours at 4°C, cryoprotected overnight in 30% sucrose, flash-frozen in hexane, and stored at −80°C. Carcasses were stored at −20°C for subsequent body composition analysis.

### 2.12. MBH micropunching and qPCR analysis

FATP1, FATP2 and FATP4 mRNA expression was quantified in a pilot cohort three weeks after FATP4-KO stereotaxic surgery. Brains were rapidly collected, frozen on dry ice and cryosectioned to the levels of the MBH [31]. MBH tissue was isolated using a sterile 2-mm biopsy punch, snap-frozen and stored at −80°C until further processing. mRNA isolation, reverse transcription and qPCR were performed as described below.

### 2.13. Brains sectioning, immunohistochemistry and image analysis

Brains were serially sectioned coronally at 20μm thickness using a cryostat (Leica Biosystems CM3050S, Germany). Sections containing the MBH were mounted on superfrost plus slides (Thermo Fisher Scientific, Switzerland) and stored in cryoprotectant (50% PBS, 30% ethylene glycol, 20% glycerol) at −20°C and processed for immunohistochemistry as described below.

#### EGFP and GFAP immunostaining

Brain sections were washed in PBS, blocked for 1 hour at room temperature in PBS containing 3% normal donkey serum and 1% bovine serum albumin (BSA) and incubated for 48 hours at 4°C with primary antibodies against GFP (1:500; Abcam) and GFAP (1:1000; Abcam). Sections were then washed and incubated for 2 hours at room temperature with Cy3 or Alexa-488 fluorophore-conjugated secondary antibodies (1:100). Complete list of primary and secondary antibodies is provided in the Supplementary Materials (Table S3). After washing, sections were counterstained with DAPI, coverslipped with Vectashield mounting medium (Nr. H-1400; Vectorlabs, Burlin-game, USA), and stored at 4°C until imaging.

#### Widefield microscopy imaging

Sections were imaged using a widefield microscope (Zeiss Axio Scan.Z1; Zeiss) with a 20× objective. Fluorescent images were analyzed using QuPath software to assess signal colocalization. At least two anatomically matched ARH or VMH sections per animal were analyzed.

### 2.14. Serum and liver immunoassay analysis

Plasma and liver triglyceride concentrations were determined using a colorimetric assay kit (Nr. 10010303; Cayman chemicals, USA). Liver triglyceride content was normalized to tissue weight and expressed as mg/g liver tissue.

Plasma leptin and insulin concentrations were measured using a U-PLEX Custom Metabolic Group 1 Kit (Nr. K15302K; Meso Scale Diagnostics, USA). Samples and standards were analyzed in duplicates, and plates were read using MESO QuickPlex SQ120 MM reader (Meso Scale Diagnostics, USA).

### 2.15. Brown adipose tissue protein extraction and Western blotting

Total protein was extracted from approximately 100 mg of interscapular brown adipose tissue (iBAT) using a lysis buffer containing protease and phosphatase inhibitors. Homogenates were clarified by centrifugation to remove lipids, and protein concentration was determined using the Pierce BCA Protein Assay Kit (Themo Fisher Scientific). Protein samples were denaturated in Laemmli buffer before electrophoresis.

Equal amounts of protein (20 μg) were separated by SDS-PAGE on 10% precast gels (Bio-Rad) and transferred to PVDF membranes, which were then blocked and incubated overnight at 4°C with primary antibodies against UCP1 (1:1000; Nr. PA124894; Thermo Fisher Scientific) and β-actin (1:5000; Nr. A5441; Sigma-Aldrich), followed by HRP-conjugated secondary antibodies. Immunoreactive bands were visualized by enhanced chemiluminescence and quantified using Fiji (ImageJ).

### 2.16. iBAT gene expression

Total RNA was extracted from iBAT using the RNeasy Lipid Tissue Mini Kit (Nr. 74804; Qiagen). cDNA was synthesized using the SensiFAST cDNA Synthesis Kit (Nr. BIO65054; Meridian Bioscience) and gene expression was quantified by SYBR Green-based quantitative PCR. UCP1 and additional thermogenic markers were analyzed using gene-specific primers. Detailed primer information is provided in the Supplementary Materials (Table S4).

### 2.17. iBAT histology

To assess the effects of MBH astrocytic FATP4 deletion and HMGCL knockdown on iBAT morphology, a portion of iBAT from mice and rats exposed to 10°C was fixed in formalin, paraffin-embedded, and sectioned at 5 μm. Haematoxylin and eosin staining was performed by the Laboratory of Animal Model Pathology (University of Zurich). Whole-slide images were analyzed using Visiopharm software to quantify adipocyte number, membrane area, and vacuole surface area.

### 2.18. Quantification and statistical analysis

Data are presented as mean ± SEM unless otherwise indicated. Statistical analyses were performed using GraphPad Prism (Version 10; GraphPad Software). Comparisons between two groups were performed using unpaired two-tailed Student’s t-tests, whereas comparisons involving two independent variables were analyzed by two-way ANOVA followed by appropriate post hoc multiple-comparison test, as specified in the figure legends. Statistical significance is denoted by \**p* < 0.05, \*\**p* < 0.01, \*\*\**p* < 0.001, \*\*\*\**p* < 0.0001

## 3. Results

### 3.1. FATP4 is the main FA transporter in mouse MBH astrocytes

To assess how MBH astrocyte lipid metabolism and signaling influence control of energy balance, we first examined the consequences of disrupting FATP4-mediated FA uptake in astrocytes of this region. The choice of the FATP transporter knockout was based on qPCR analyses of primary astrocytes and neurons derived from the MBH of mice, which identified FATP4 as the main long-chain FA transporter in the MBH (**Fig. 1A**). FATP4 expression was astrocyte-specific and significantly higher than that of other FATP isoforms (**Fig. 1A**).

**Figure 1.**
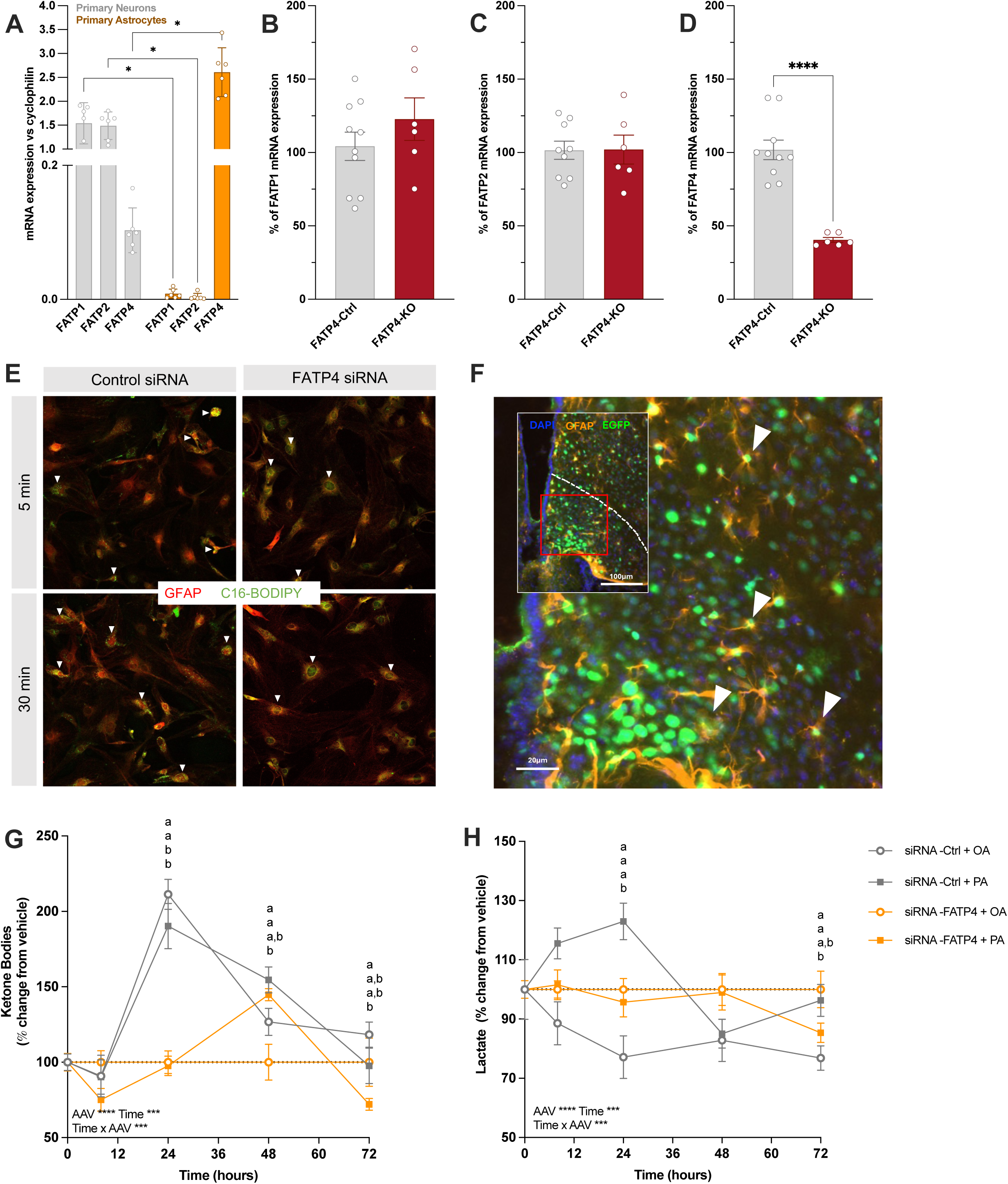
FATP4 expression in primary hypothalamic cells and following its deletion in astrocytes of the MBH. (A) Gene expression analysis of FATP isoforms in primary neurons vs. primary astrocytes isolated from the MBH of wild-type mice. (B-D) qPCR quantification of MBH micropunches from FATP4-Ctrl and -KO mice showing the expression of (B) FATP1, (C) FATP2, (D) FATP4. n=5-10 per group. Data are presented as a percentage of mRNA expression relative to the control group, with two housekeeping genes used for normalization. (E) FATP4 siRNA treated primary hypothalamic astrocytes (right) and control cells (left) after 5 and 30 minutes of incubation with 10 μM GFP-BODIPY-PA. PA is visualized in green, GFAP in red. Arrows indicate astrocytes and BODIPY-PA colocalization. (F) Representative immunofluorescent images of MBH sections from FATP4-KO mice showing DAPI (blue), AAV EGFP (green), and GFAP (orange). White arrows indicate colocalization of AAV EGFP with the astrocytic marker GFAP, confirming astrocyte-specific viral targeting. Primary MBH astrocyte ketone body (G) and lactate (H) production in response to oleic acid (OA; 20 μM) and palmitic acid (PA; 20 μM). Data are expressed as percentage change relative to vehicle treatment. Data are presented as mean ± SEM and analyzed using unpaired t-test (A-D), repeated-measure two-way ANOVA (G-H). \**p* < 0.05, \*\**p* < 0.01, \*\*\**p* < 0.001, \*\*\*\**p* < 0.0001. ^a, b^Data points with differing superscripts differ from each other at p < 0.05.

To determine whether FATP4 reduces astrocytic palmitic acid (PA) uptake, we performed a fluorescent FA assay using the palmitate analogue BODIPY-PA (10 μM). This probe was selected because PA is the predominant saturated FA present in the cerebrospinal fluid [32], allowing us to assess the impact of FATP4 knockdown on the uptake of a physiologically relevant FA. First, qPCR confirmed an 89% reduction in Slc27a4 (FATP4) mRNA levels in FATP4-siRNA cells. Additionally, the knockdown significantly reduced the update of BODIPY-PA by primary hypothalamic astrocytes. Qualitative assessment revealed lower intracellular accumulation of the fluorescent PA analogue in FATP4-siRNA-treated astrocytes compared to control cells after both 5 and 30 minutes of incubation. Quantification of BODIPY-PA colocalization with GFAP demonstrated a reduction from 25-45% after 5 minutes, and from 22-51% after 30 minutes in FATP4-depleted cells relative to controls (**Fig. 1E**).

### 3.2. Astrocyte fatty acid uptake is necessary for FA-induced ketone production

Having established efficient FATP4 knockdown, we next examined its impact on astrocyte ketone body and lactate production. Levels for these two metabolites were measured in the media of hypothalamic astrocytes following FATP4-siRNA treatment and oleic acid (OA) or PA exposure.

FATP4 knockdown significantly reduced ketone body production. When normalized to vehicle-treated cells, FATP4-depleted astrocytes showed a blunted ketogenic response to both OA and PA at 24 hours (+11% and +8%, respectively) compared with control-siRNA treated cells (+111% and 90%, respectively). After 72 hours of FA exposure, ketone body levels returned to baseline in all groups (**Fig. 1G**). Interestingly, OA but not PA treatment increased lactate production in FATP4-siRNA astrocytes at 24 and 72 hours (**Fig. 1H**). Together, these findings indicate that FATP4 depletion shifts astrocyte FA metabolism away from ketogenesis and potentially towards lactate production, particularly in response to OA.

### 3.3. Validation of the FATP4 depletion in MBH GFAP-positive astrocytes

We used a viral-mediated Cre/lox recombination strategy to selectively target the FATP4 gene, administering a Cre-expressing AAV directly into the MBH of FATP4^flox/flox^ mice. Cre recombinase was expressed under the control of the human GFAP (hGFAP) promoter, a well-established astrocyte marker. The fluorescent reporter EGFP was also driven by hGFAP, enabling the identification of transfected, Cre-expressing astrocytes. In mice injected with the Cre-containing AAV (Cre^MBH-hGFAP^: FATP4^flox^; FATP4-KO), EGFP signals were confined to the nucleus, consistent with the nuclear localization of the Cre recombinase (**Fig. 1F**). Immunohistochemical analysis of MBH-containing slices was used to verify the co-localization of EGFP with GFAP (**Fig. 1F**). In parallel, MBH micropunches were then analyzed to quantify the degree of FATP4 knockout. As expected, the Cre-AAV reduced FATP4 expression by approximately 60% relative to controls (**Fig. 1D**). Because FATP4 is predominantly expressed in astrocytes, this reduction was specific to astroyctes. Notably, no compensatory increases in the expression of other FATP isoforms were detected (**Fig. 1B-C**). **Fig. 2A** outlines the experimental timeline for the FATP4 study. Following stereotaxic surgery, mice were maintained on a chow diet for two weeks. Body composition analysis confirmed that initial fat and lean mass were comparable between groups (data not shown). Body weight and food intake during this period were also very similar (data not shown).

**Figure 2.**
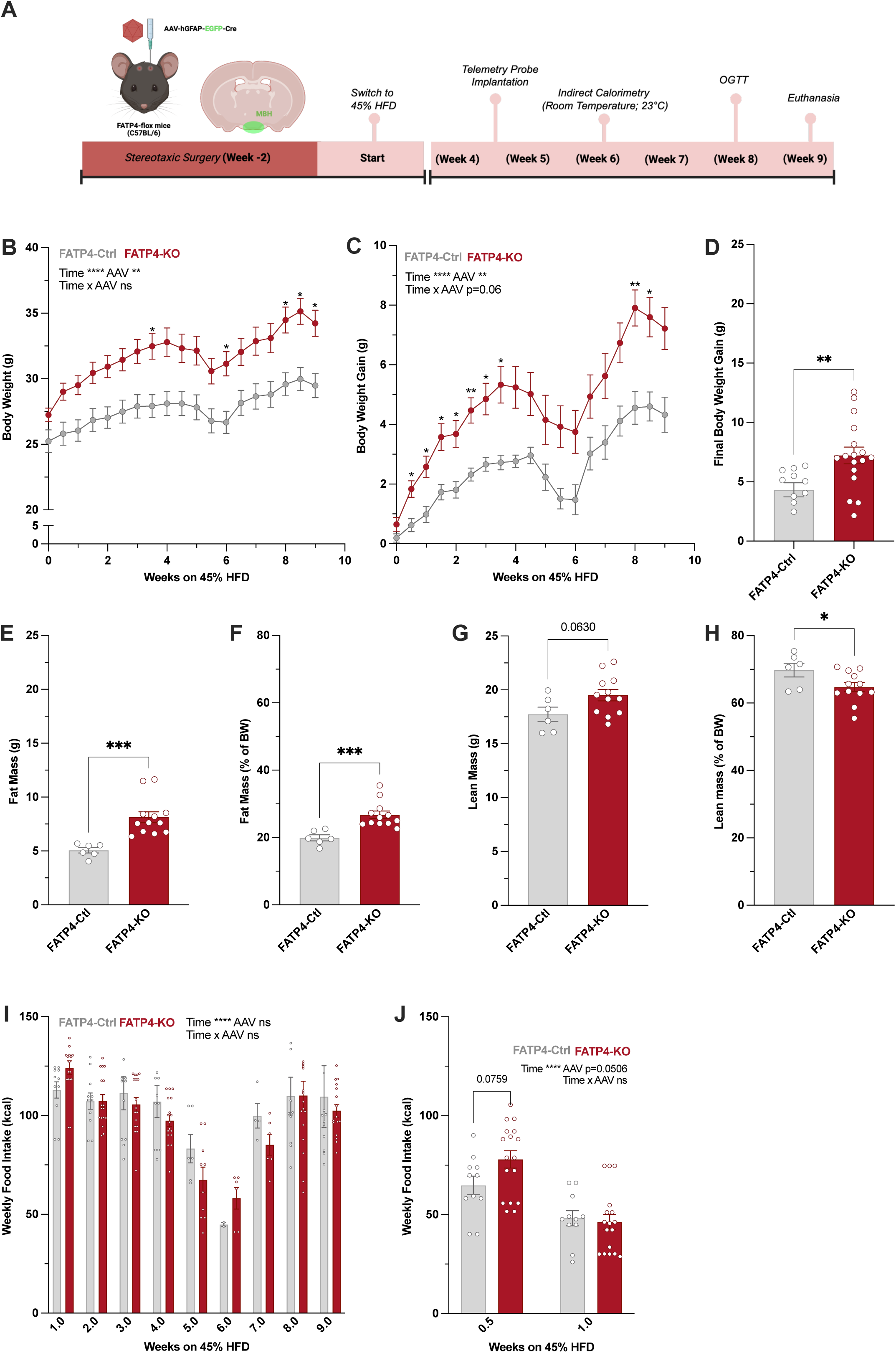
Astrocyte-specific deletion of FATP4 in the MBH altered HFD-induced body weight gain and fat accumulation. (A) Experimental timeline following stereotaxic surgery for astrocyte-specific FATP4 knockout in the MBH. (B) Absolute body weight and (C) weight gain over the course of the 45% HFD exposure. (D) Final body weight gain. (E-H) Body composition analysis at the end of the study: fat mass (E-F) and lean mass (G-H) presented as absolute mass and normalized to terminal body weight. (I) Weekly food intake throughout the experimental period. (J) Food intake during the first week of HFD is split into two intervals: 0.5 and 1. n=6-17 per group. Data are presented as mean ± SEM and analyzed using repeated-measure two-way ANOVA or unpaired t-test as appropriate. *p < 0.05, **p < 0.01, ***p < 0.001, ****p < 0.0001.

### 3.4. MBH astrocyte-specific FATP4 knockout exacerbates diet-induced obesity by reducing energy expenditure

Owing to their resemblance to the average Western diet, 45% HFD have been widely used to investigate the development of metabolic disorders such as obesity and type 2 diabetes [33]. Exposure to this diet triggered a steady and progressive increase in body weight in both groups (**Fig. 2B**). FATP4-KO mice displayed a significantly greater weight gain than their FATP4-Ctrl counterpart, indicating an enhanced susceptibility to diet-induced obesity (p<0.01; **Fig. 2C-D**). This phenotype was further highlighted by body composition analyses, which revealed a ∼61% increase in absolute fat mass and a 35% increase in fat mass expressed as a proportion of total body weight in knockout animals compared to controls (p<0.001; **Fig. 2E-F**). In contrast, lean mass remained unchanged or proportionally reduced (**Fig. 2G-H**).

Energy homeostasis results from the dynamic balance between energy intake and energy expenditure. During the first week on HFD, we detected a modest but short-lived rise in caloric intake in knockout mice (**Fig. 2I**). Studies in DIO and DR rats have shown that the initial week of HFD exposure is critical, with DR rats requiring approximately three days before they can reduce their food intake (a process linked to MBH ketogenesis) [15]. Motivated by these observations, we broke down our first week of HFD into two timepoints (week 0.5 and week 1; **Fig. 2J**). This analysis demonstrated that a slight increase in caloric intake was primarily driven by alterations at week 0.5, where FATP4-KO mice showed a trend towards higher food consumption (∼20% increase; p=0.0759). After this early divergence, weekly food intake remained comparable between the two groups for the remainder of the study (**Fig. 2I**).

Indirect calorimetry was used to assess the impact of MBH astrocyte FATP4 ablation on energy expenditure (EE) and respiratory exchange ratio (RER). To capture potential temporal shifts in these metabolic parameters, independent cohorts of mice were placed in the calorimetry cages at different timepoints throughout the study. EE and RER were first assessed during the initial week of 45% HFD exposure. During this early phase, no differences in EE or RER were observed between FATP4-KO mice and controls during either the light or dark phases (**Fig. 3A-E**).

**Figure 3.**
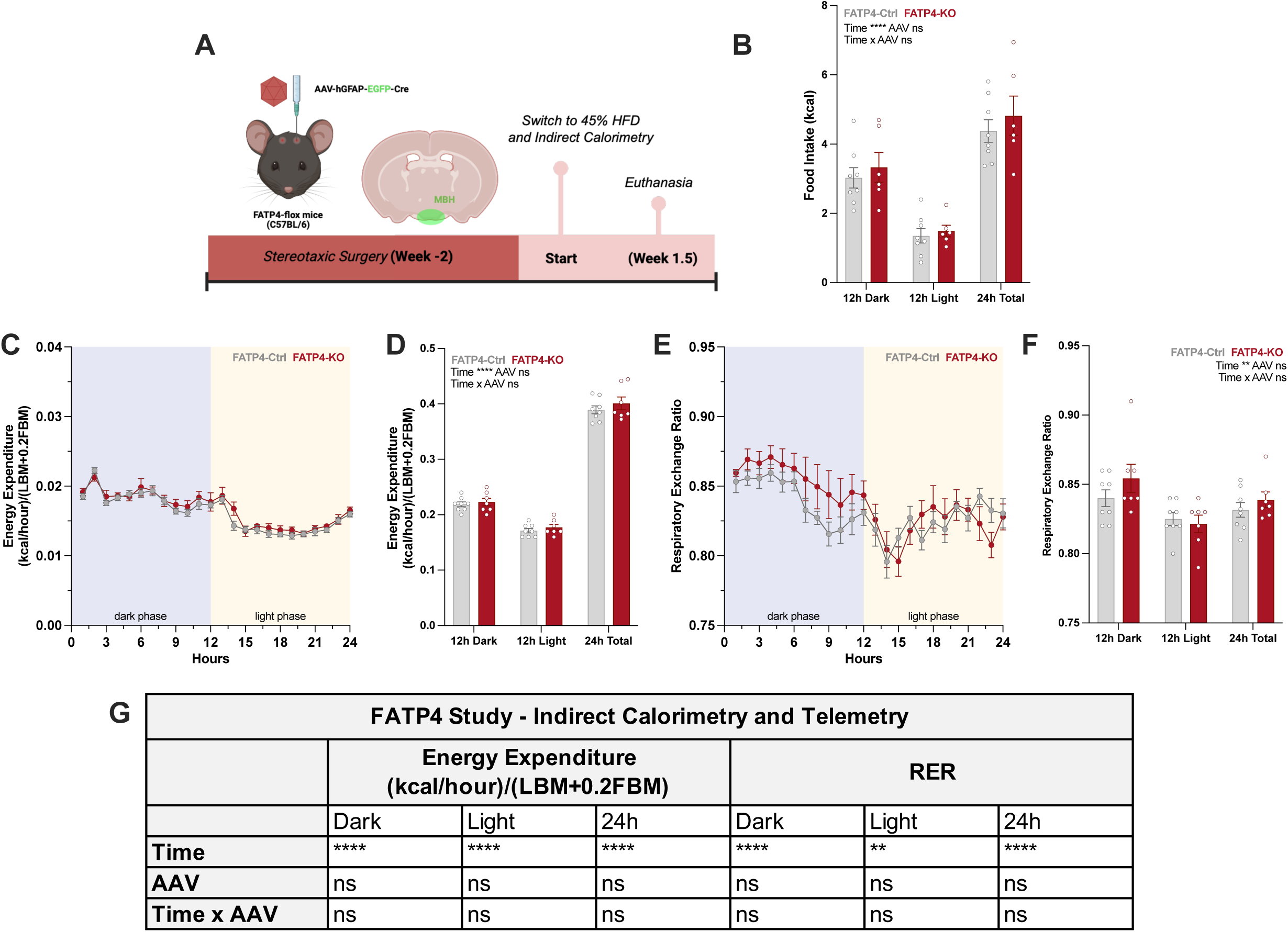
Effects of FATP4-KO on energy expenditure and substrate utilization during early HFD exposure. (A) Experimental timeline. (B) Three-day average food intake across dark- and light-phases. (C) Three-day average energy expenditure (EE) normalized to terminal body composition. Line plots depict hourly profiles, with dark and light phases in blue and yellow, respectively. (D) 12-hour phase-specific average EE. (E-F) Respiratory exchange ratio (RER) presented as (E) three-day average hourly profiles and (F) phase-averaged values. (G) Summary statistics of indirect calorimetry measurements corresponding to panels C and E, with analyses conducted on dark phase, light phase and 24-hour total values. n=7-8 per group. Data are presented as mean ± SEM and analyzed using repeated-measure two-way ANOVA. \**p* < 0.05, \*\**p* < 0.01, \*\*\**p* < 0.001, \*\*\*\**p* < 0.0001.

In contrast, after 5 weeks on 45% HFD, FATP4-KO mice consistently exhibited lower EE compared with FATP4-Ctrls (**Fig. 4A**). Quantitative analyses of total, dark-phase and light-phase EE confirmed a significant overall decrease in the KO group (**Fig. 4B**). The latter changes occurred in the absence of any differences in total activity (data not shown), indicating that the reduction in EE was not secondary to decreased locomotion. In parallel, hourly analysis of RER did not reveal a major overall effect of the knockout on substrate utilization across the 24-hour cycle (**Fig. 4C-D**). Both FATP4-Ctrl and -KO mice exhibited similar circadian oscillations, with RER values rising during the active (dark) phase (reflecting increased carbohydrate utilization) and declining during the resting (light) phase, consistent with lipid oxidation (**Fig. 4C**). Although a visible decrease in RER could be observed in FATP4-KO mice during the light phase in the hourly trace (**Fig. 4C**), these differences did not reach statistical significance at individual timepoints. Nonetheless, when RER values were averaged over the entire 12-hour light period, a significant AAV-dependent reduction emerged (**Fig. 4D**), resulting in a significant interaction between time and AAV. These results indicate an increased reliance on lipid oxidation in knockout animals during the light phase. In addition to EE and RER, the indirect calorimetry system allowed monitoring of food intake. Consistent with weekly food intake measurements, no statistical differences were observed in 12-h averaged food intake between groups (**Fig. 4E**).

**Figure 4.**
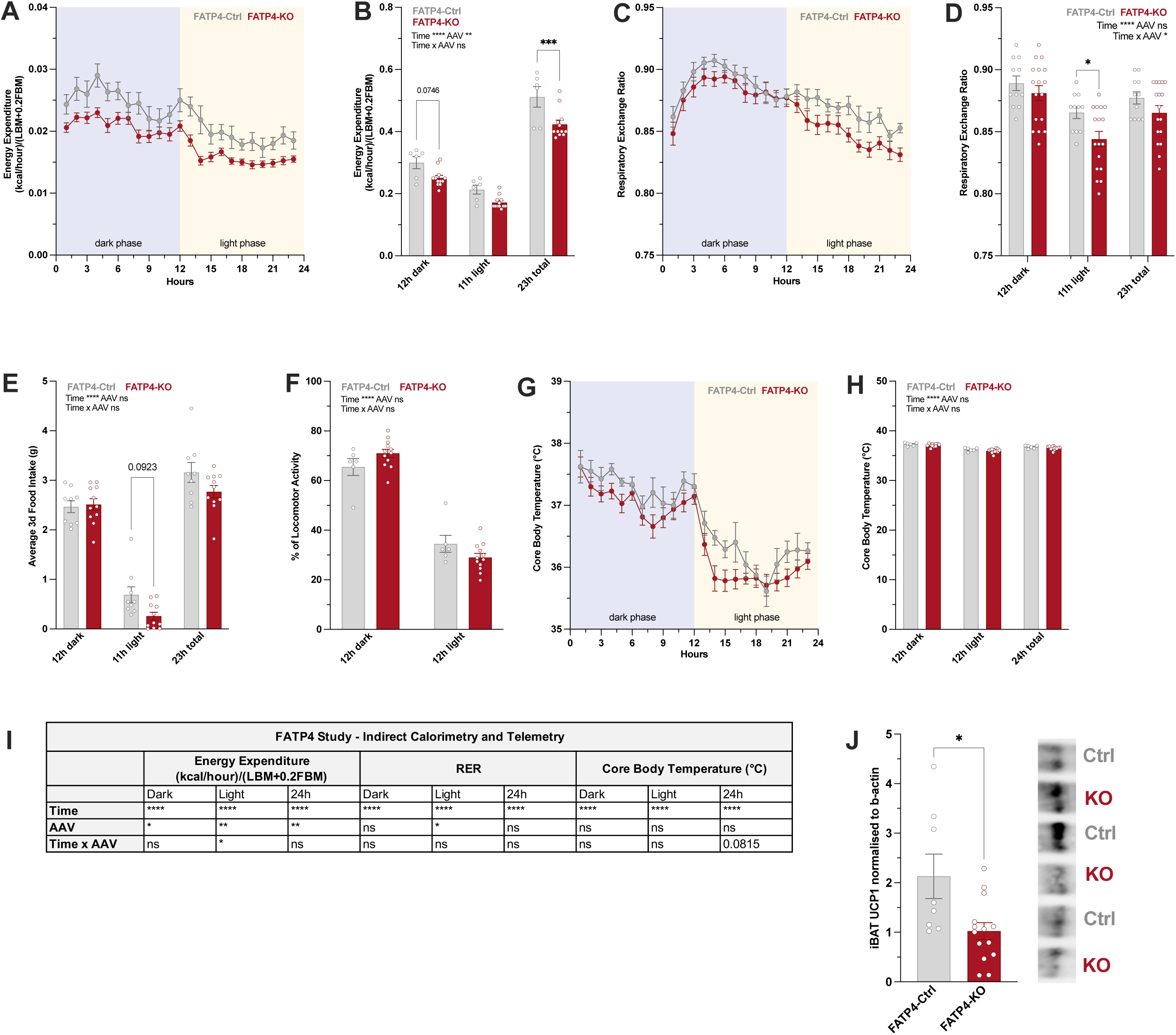
Effects of FATP4-KO on energy expenditure and substrate utilization after 5 weeks of HFD. (A) Three-day average energy expenditure normalized to body composition data. Line plots depict hourly profiles, with dark and light phases in blue and yellow, respectively. (B) 12h-phase-specific average EE. (C-D) Respiratory exchange ratio (RER) presented as (C) three-day average hourly profiles and (D) phase-averaged values. (E) Phase-averaged food intake during the 3-day indirect calorimetry recordings. (F) Locomotor activity. (G-H) Core body temperature was further analyzed as (G) hourly and (H) phase-averaged values. (I) Summary statistics of indirect calorimetry and telemetry measurements corresponding to panels A, C and G, with analyses conducted on dark phase, light phase and 24-hour total values. (J) Western blot analysis of UCP1 protein levels in the iBAT, normalized to β-actin. n=6-17 per group. Data are presented as mean ± SEM and analyzed using repeated measure two-way ANOVA followed by post hoc testing, or unpaired t-test. \**p* < 0.05, \*\**p* < 0.01, \*\*\**p* < 0.001, \*\*\*\**p* < 0.0001.

### 3.5. MBH astrocyte FATP4 loss reduces iBAT UCP1 expression without altering core body temperature

Both locomotor activity and core body temperature are recognized determinants of total EE [34]. To assess whether FATP4 ablation affected either of these measures, mice were implanted with intraperitoneal telemetry sensors. Locomotor activity did not differ between FATP4-KO and FATP4-Ctrl mice during either the dark or light phases (**Fig. 4F**). While no significant effect of the AAV on hourly core body temperature was detected (**Fig. 4G**), a trend (p=0.0815) towards a lower body temperature during the light phase was observed (**Fig. 4G, I**). Core body temperature was likewise comparable between groups when averaged across each 12-hour phase (**Fig. 4H**). Brown adipose tissue (BAT) is a primary site of adaptive heat production, driven by the activity of mitochondrial uncoupling protein 1 (UCP1). Western blot analysis of interscapular BAT (iBAT) demonstrated a significant reduction in UCP1 protein levels in FATP4-KO mice relative to controls (**Fig. 4J**). While decreased UCP1 expression is likely to compromise iBAT thermogenic capacity, this alteration alone may not be sufficient to impact whole-body core temperature.

### 3.6. MBH astrocyte FATP4 knockout leads to early disruption of glucose homeostasis

To further characterize the role of MBH astrocytic FATP4 in peripheral metabolism, we performed an oral glucose tolerance test after eight weeks on 45% HFD. At baseline, FATP4-KO mice already exhibited elevated fasting glucose levels (**Fig. 5A**). Following the administration of a glucose bolus, FATP4-Ctrl mice cleared circulating glucose more efficiently than knockouts, which showed higher glucose levels at 15 minutes post-challenge, and a significantly greater area under the curve (AUC), calculated from the baseline level of each mouse (**Fig. 5B**), indicative of impaired glucose tolerance. We also examined the effects of FATP4 ablation on circulating insulin and leptin levels; however, neither of these parameters differed between groups (**Fig. 5C-D**) despite the increase in fat mass. Overall, the combination of higher fasting glucose and normal insulin levels suggests an early disruption of systemic glucose homeostasis in FATP4-KO mice, correlating with the increased body weight.

**Figure 5.**
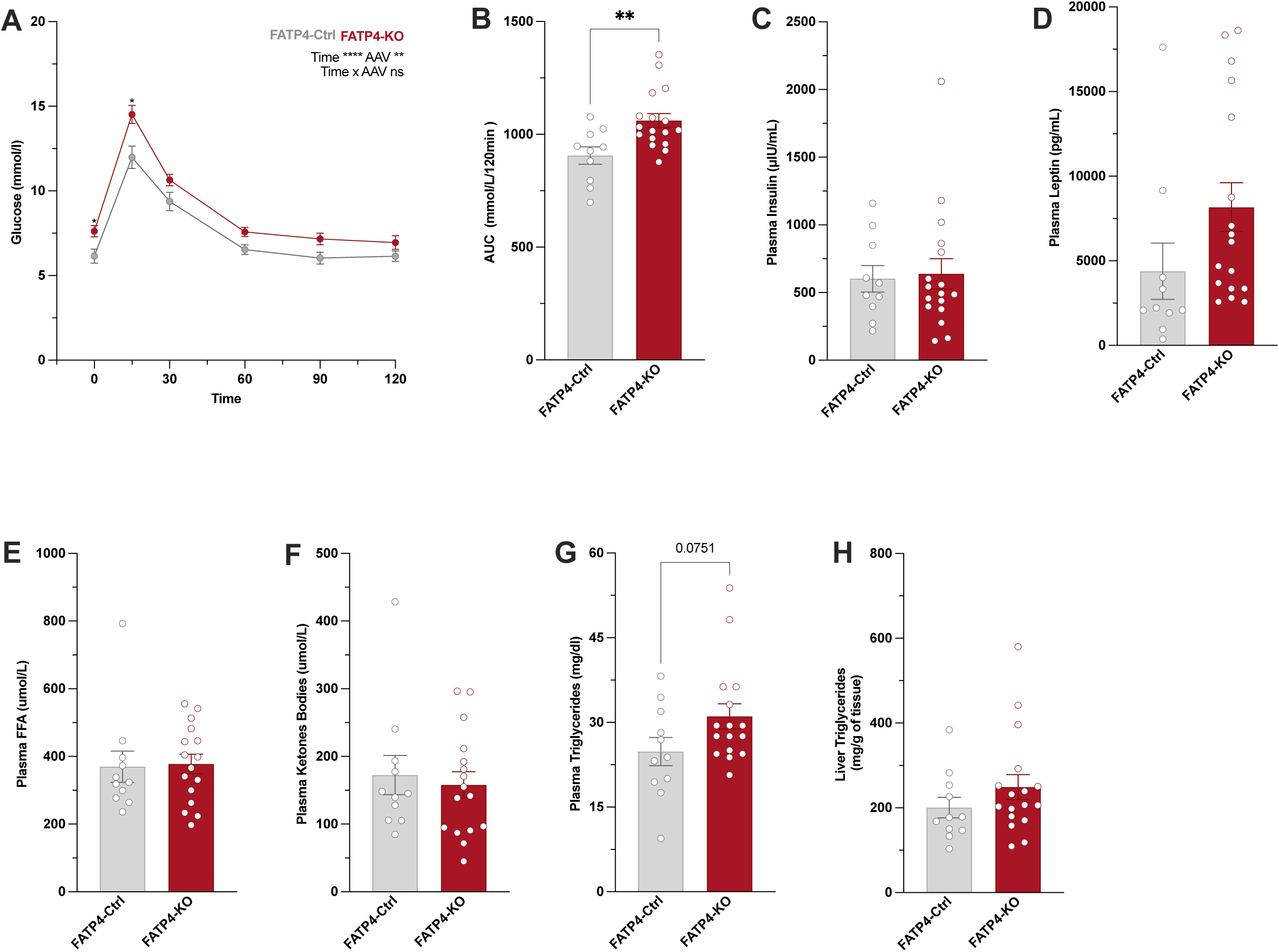
MBH astrocyte-specific deletion of FATP4 affected peripheral glucose levels. (A) Oral glucose tolerance test showing blood glucose levels over time following glucose administration in FATP4-Ctrl and FATP4-KO mice. (B) Area under the curve (AUC) analysis of the OGTT. Plasma (C) insulin, (D) leptin, (E) free fatty acids, and (F) ketone body levels. (G) Plasma triglyceride and (H) liver triglyceride levels. Except for glucose measurements, all plasma and liver samples were collected during euthanasia and processed *ex vivo*. n=10-17 per group. Data are presented as mean ± SEM. OGTT time-course data were performed using a two-way ANOVA with multiple comparisons; all other parameters were analyzed using unpaired t-test. \**p* < 0.05, \*\**p* < 0.01, \*\*\**p* < 0.001, \*\*\*\**p* < 0.0001.

Plasma free fatty acids (FFA) and ketone body levels were comparable between groups, indicating that FATP4 deletion did not affect hepatic FFAs mobilization or peripheral ketogenesis (**Fig. 5E-F**). Liver triglyceride content was likewise unchanged (**Fig. 5H**). In contrast, plasma triglyceride levels tended to be higher in FATP4-KO mice than in controls (∼25%; p=0.0751), pointing to a potential alteration in systemic lipid homeostasis (**Fig. 5G**).

### 3.7. Knocking down MBH astrocyte HMGCL partially recapitulates the metabolic phenotype of FATP4 deletion

Following the latter study, we sought to determine whether selectively disrupting MBH astrocyte ketogenesis would reproduce the metabolic deficits observed in FATP4-KO mice. To this end, we targeted the expression of HMGCL, a critical component of the ketogenic pathway [35,36]. Since HMGCL^fl/fl^ mice are not available, and to test our hypothesis across species, we employed an alternative approach using Sprague-Dawley rats. Specifically, we knocked down HMGCL expression via stereotaxic delivery of an AAV encoding hGFAP-driven anti-HMGCL shRNAs into the MBH (shRNA^MBH-hGFAP^: HMGCL; hereafter referred to as HMGCL-KD). A non-targeting shRNA was used in the control AAV (shRNA^MBH-hGFAP^: Ctrl; HMGCL-Ctrl) (**Fig. 6A**). Once again, EGFP expression enabled the verification of successful MBH astrocyte transfection. Qualitative analyses of rat brain sections confirmed co-localization of the reporter with GFAP (**Fig. 6B**). Although a fraction of EGFP signals was detected in HuCD-positive neurons (**Fig. 6C**), it is important to note that neurons do not express GFAP and their ketogenic gene expression is minimal to negligible [37]. Therefore, this off-target neuronal transfection is unlikely to confound either the metabolic outcomes observed or the interpretation of our results.

**Figure 6.**
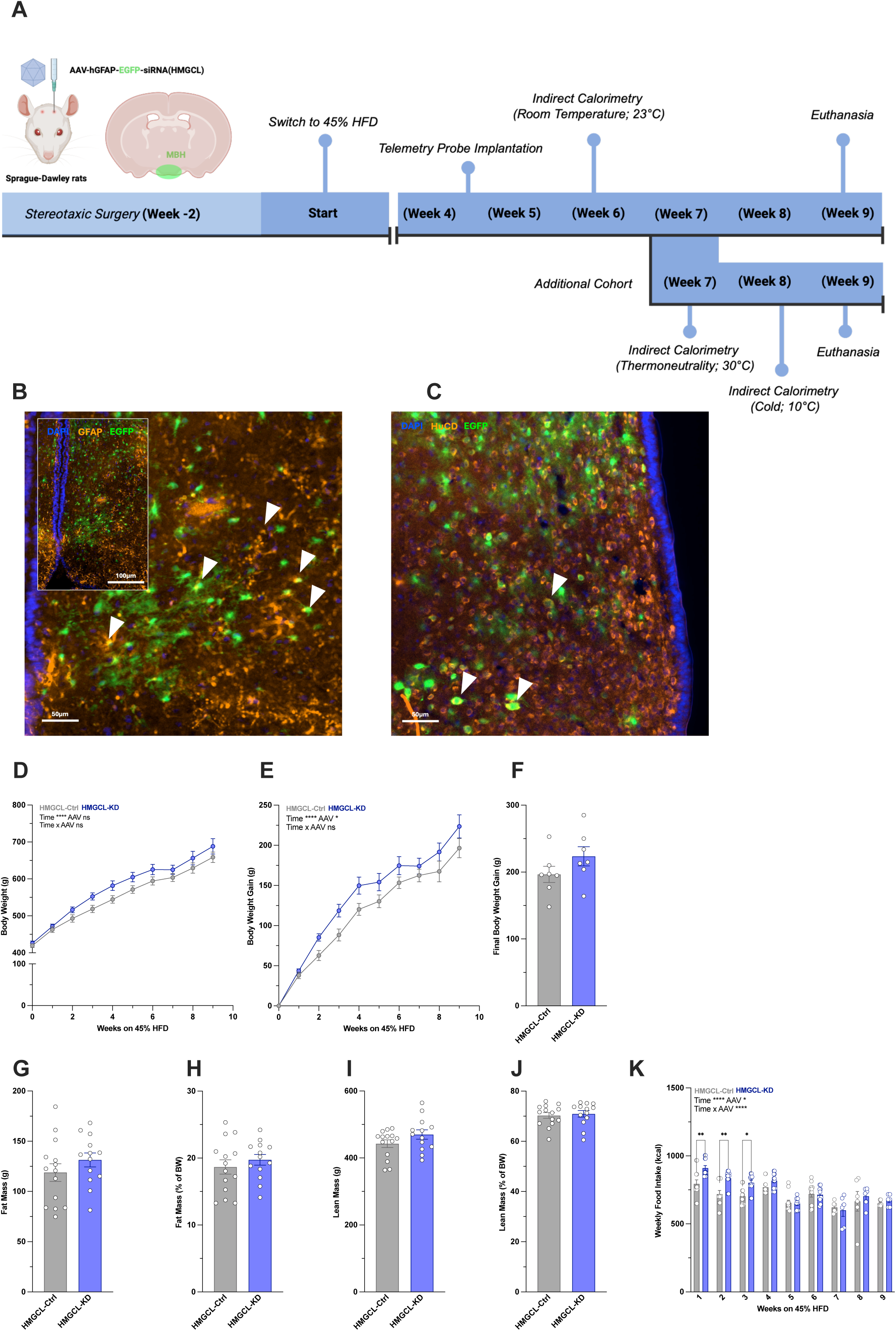
Astrocyte-specific knockdown of HMGCL in the MBH promoted HFD-induced body weight gain and transiently increased food intake. (A) Experimental timeline. (B-C) Representative immunofluorescent images of MBH sections from HMGCL -KO rats showing DAPI (blue), AAV EGFP (green), and (B) GFAP or (C) HuCD (orange). White arrows in panel A indicate colocalization of EGFP with GFAP, confirming astrocyte-specific viral targeting. In panel C, white arrows indicate colocalization of EGFP with neuronal marker HuCD, evidencing unwanted neuronal transfection. (D) Absolute body weight and (E) body weight gain over the course of 45% HFD exposure. (F) Final body weight gain was measured at week 9. (E-H) Body composition analysis at the end of the study: fat mass (G-H) and lean mass (I-J), are presented as absolute mass and normalized to terminal body weight. (K) Weekly food intake throughout the experimental period. n=6-14 per group. Data are presented as mean ± SEM and analyzed using repeated-measure two-way ANOVA or unpaired t-test as appropriate. \**p* < 0.05, \*\**p* < 0.01, \*\*\**p* < 0.001, \*\*\*\**p* < 0.0001.

Similar to the FATP4 knockout model, disrupting MBH astrocyte ketogenesis exacerbated the metabolic effects of 45% HFD exposure. HMGCL knockdown led to a greater HFD-induced weight gain, with KD rats gaining ∼13% more weight than controls (**Fig. 6D-F**). However, body composition analyses revealed no significant differences in either fat mass or lean mass between the two groups (**Fig. 6G-K**). During the first three weeks on HFD, HMGCL-KD rats displayed a transient yet significant increase in food intake (**Fig. 6K**), suggesting impaired adaptation to the diet’s increased caloric density. Thereafter, caloric intake stabilized and remained comparable to that of controls (**Fig. 6K**).

Indirect calorimetry pointed to phase-specific rather than global changes in EE. When examining EE across the full 24-hour cycle, the depletion of MBH astrocyte HMGCL had no effect (**Fig. 7A**). However, restricting the analysis to the dark phase revealed a significant reduction by 20% in EE in HMGCL-KD rats (**Fig. 7A, H**). This phase-specific effect was further supported by the 12-hour averaged data: although total, dark-phase and light-phase EE did not differ significantly between HMGCL-KD and HMGCL-Ctrl animals in post-hoc comparisons, the ANOVA revealed a significant Time x AAV interaction (**Fig. 7B**). These findings suggest that the knockdown may not alter overall daily EE, but reduce EE specifically during the dark, active phase. Hourly and 12-hour averaged RER values did not reveal statistically significant group differences (**Fig. 7C-D**).

**Figure 7.**
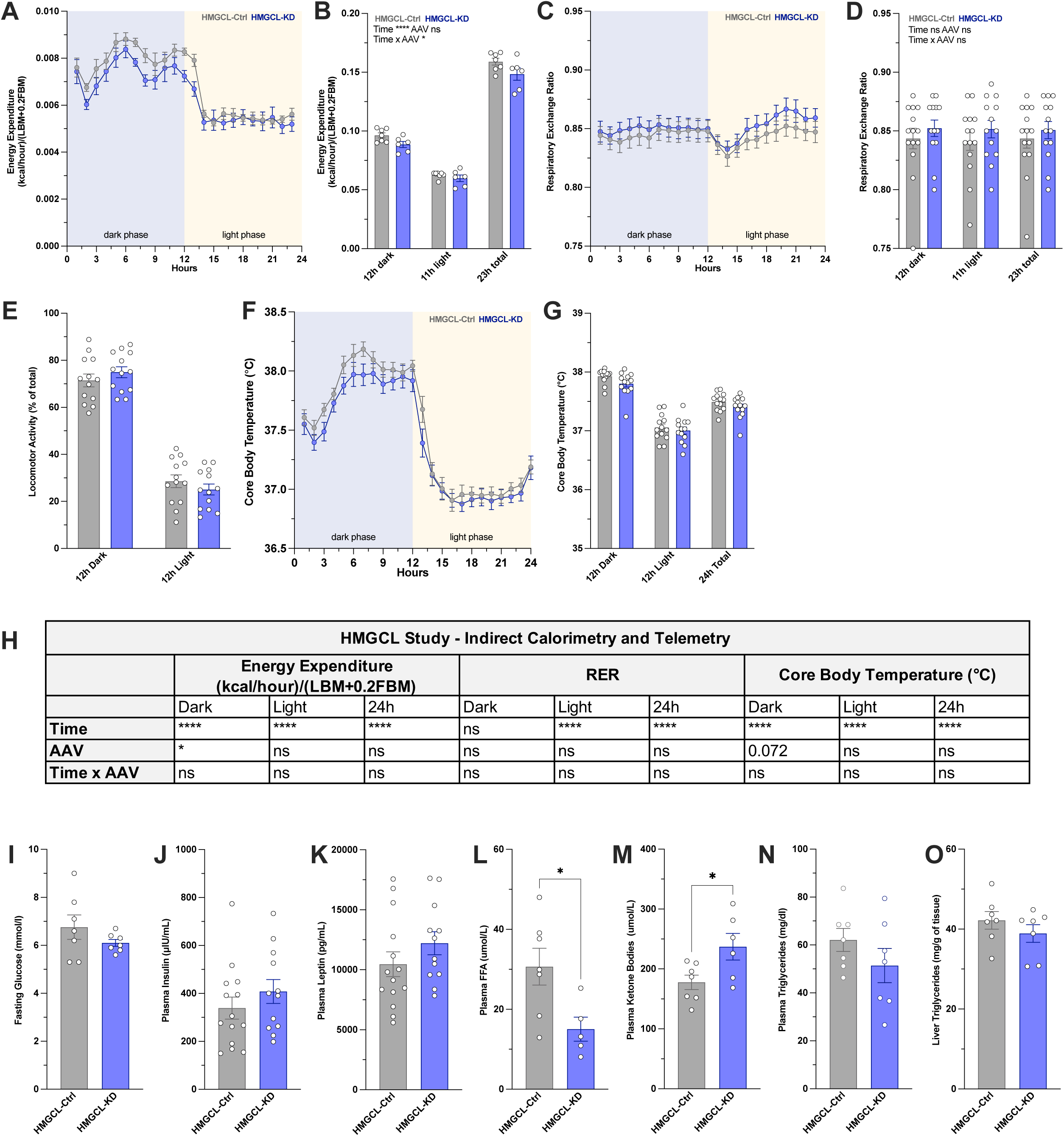
Effects of HMGCL-KD on energy expenditure and substrate utilization after 6 weeks of HFD. (A) Three-day average energy expenditure normalized to body composition data. Line plots depict hourly profiles, with dark and light phases in blue and yellow, respectively. (B) 12h-phase-specific average energy expenditure. (C-D) Respiratory exchange ratio (RER) across dark and light phases, shown as (C) hourly and (D) phase-averaged values. (E) Locomotor activity. (F-G) Core body temperature measured by continuous telemetry recordings. Body temperature was further analyzed as (F) hourly and (G) phase-averaged values. (H) Summary statistics of indirect calorimetry and telemetry measurements corresponding to panels A, C and F, with analyses conducted on dark phase, light phase and 24-hour total values. Fasting plasma (I) glucose levels. (J) insulin, (K) leptin, (L) free fatty acids, and (M) ketone body levels. (N) Plasma triglyceride and (O) liver triglyceride levels. Except for glucose measurements, all other plasma and liver parameters were processed *ex vivo*. n=9-16 per group. Data are presented as mean ± SEM and analyzed using repeated measure two-way ANOVA followed by post hoc testing or unpaired t-test as appropriate. *p < 0.05, **p < 0.01, ***p < 0.001, ****p < 0.0001.

Locomotor activity also remained unchanged between the two groups (**Fig. 7E**). Although phase-averaged core body temperatures were similar, hourly analysis suggested a mild downward shift in HMGCL-KD rats during their active period, consistent with a trend (p =0.072) toward reduced temperature (**Fig. 7F-H**) that may contribute to the reduced energy expenditure observed at this time.

### 3.8. MBH astrocyte HMGCL knockdown preserves glucose homeostasis but alters lipid metabolism

In contrast to FATP4-KO mice, we did not detect significant differences in fasting plasma glucose, insulin or leptin levels (**Fig. 7I-J**). Plasma and liver triglyceride levels also did not differ (**Fig. 7N, O**). Conversely, plasma FFA levels (**Fig. 7L**) were significantly reduced in HMGCL-KD rats compared to controls, whereas circulating ketone levels were significantly increased (**Fig. 7M**). Together, these findings indicate that while peripheral glucose homeostasis may be preserved, specific aspects of peripheral lipid metabolism may become altered upon disruption of MBH astrocyte ketogenesis.

### 3.9. MBH astrocyte knockdown affects iBAT thermogenic adaptation to cold challenge

To further assess the role of MBH astrocyte-derived ketone body signaling in thermogenesis, we examined a separate cohort of rats housed in a climate chamber, enabling simultaneous calorimetry and telemetry measurements under controlled thermal conditions.

At thermoneutrality (30°C), EE and RER values of control and knockdown rats closely overlapped across the 24-hourly recordings (**Fig. 8A-D**). Under these conditions, locomotor activity and core body temperature were also comparable across groups (**Fig. 8E-G**).

**Figure 8.**
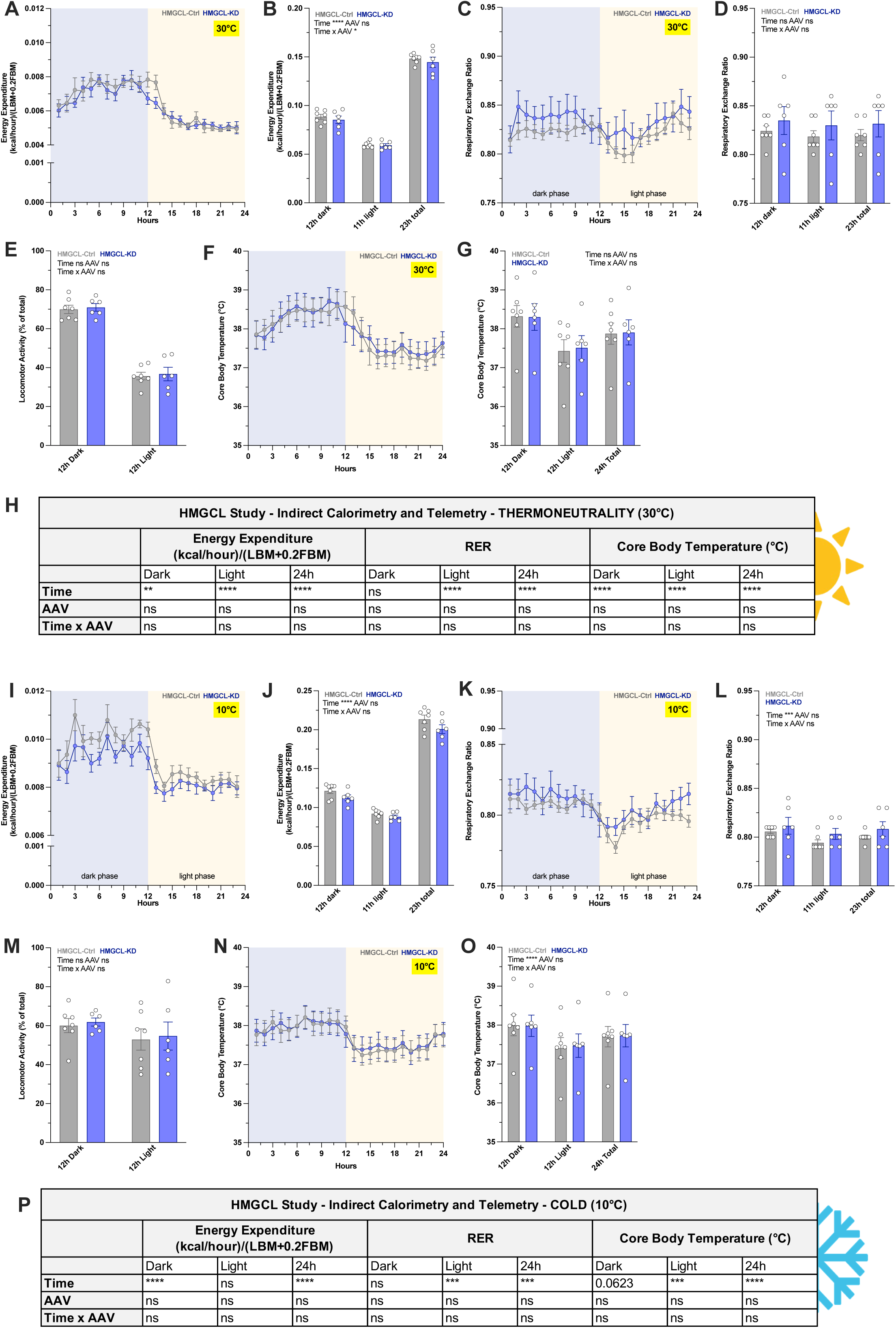
Effects of MBH astrocyte HMGCL-KD on energy expenditure and substrate utilization at thermoneutrality and during cold exposure after 7 weeks of HFD. (A-H) Recordings at thermoneutral conditions (30 °C) or (I-O) at cold conditions (10 °C). (A, I) Three-day average energy expenditure normalized to body composition data. Line plots depict hourly profiles, with dark and light phases in blue and yellow, respectively. (B, J) 12h-phase-specific average energy expenditure. (C-D, K-L) Respiratory exchange ratio across dark and light phases, shown as (C, K) hourly and (D, L) phase-averaged values. (E, M) Locomotor activity. (F-G, N-O) Core body temperature measured by continuous telemetry recordings. (H, P) Summary statistics of indirect calorimetry and telemetry measurements corresponding to panels A, C and F (30 °C) and I, K, and N (10 °C), with analyses conducted on dark phase, light phase and 24-hour total values. n=6-7 per group. Data are presented as mean ± SEM and analyzed using repeated measure two-way ANOVA followed by post hoc testing. *p < 0.05, **p < 0.01, ***p < 0.001, ****p < 0.0001.

Acute exposure to cold (10°C) elicited the expected increase in total EE compared to room temperature and thermoneutrality in both groups (**Fig. 8I-J**). No statistically significant differences in EE or RER were detected under these conditions (**Fig. 8I-L**). Cold exposure also led to a reduction in locomotor activity during the dark phase in the two groups (**Fig. 8M**), but core body temperature remained remarkably stable and indistinguishable across HMGCL-KD and HMGCL-Ctrl animals (**Fig. 8N-O**).

Both experimental groups were euthanized under ongoing cold exposure, and iBAT samples were collected for *ex vivo* analysis. First, the expression of β3AR and UCP1 was decreased by ∼70% (P=0.08) and ∼40% (P=0.12), respectively, in HMGCL-KD rats compared to control rats (**Fig. 9A-B**). β3AR is the primary adrenergic receptor in the BAT, and as such, lower expression may reflect decreased sympathetic output to this tissue, whereas reduced UCP1 expression could indicate impaired thermogenic capacity [38,39]. These transcriptional patterns aligned with the obtained histological findings: HE staining showed that although the number of iBAT adipocytes was comparable across groups, knockdown animals displayed altered cellular morphology, characterized by a significantly smaller cytoplasmic area and an increase in lipid vacuolar area compared to controls (**Fig. 9C-E**). Given the cold conditions used in this experiment, these differences may suggest that control animals underwent a more effective, cold-induced browning of iBAT than HMGCL-KD rats. Moreover, these effects may reflect a subtle influence of MBH astrocyte function on sympathetic nervous system activity in rats.

**Figure 9.**
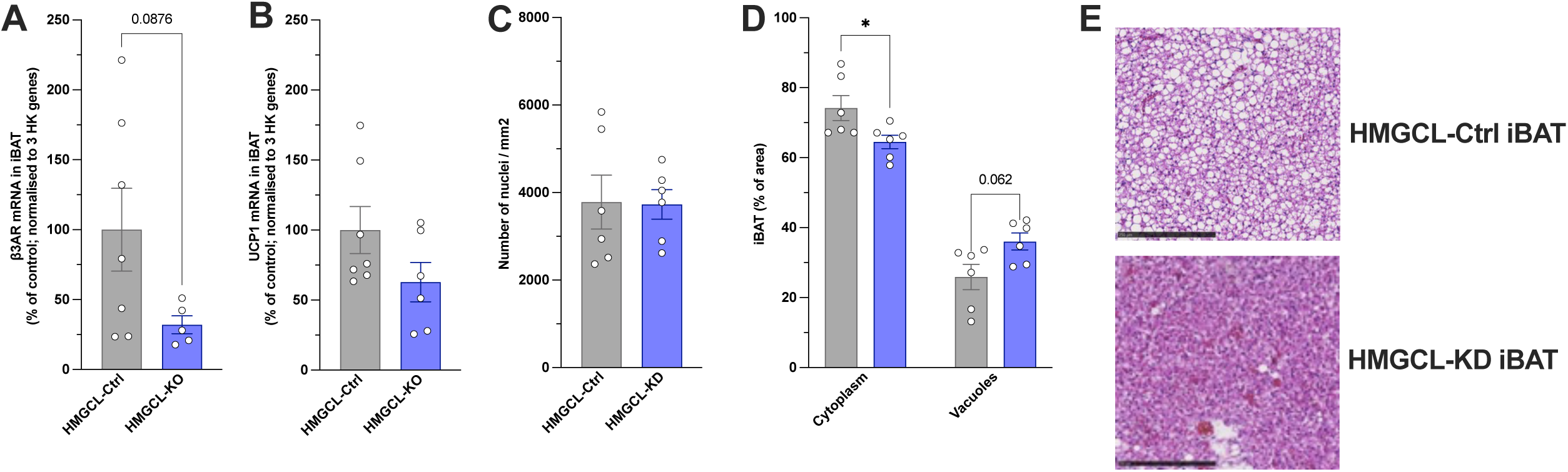
HMGCL-KD altered iBAT gene expression and adipocyte morphology. qPCR analysis of (A) β3-AR and (B) UCP1 mRNA levels relative to controls. Data are presented as a percentage of mRNA expression relative to the control group, with three housekeeping genes used for normalization. (C-E) HE staining iBAT analysis: (C) Quantification of the number of nuclei per mm². (D) Quantification of cytoplasmic and vacuolar area, expressed as a percentage of total tissue area. (E) Representative HE–stained images from iBAT of HMGCL-Ctrl and HMGCL-KD rats (scale bar = 250 µm). n=6-7 per group. Data are presented as mean ± SEM and analyzed using unpaired t-tests, or multiple t-tests for panel D. *p < 0.05, **p < 0.01, ***p < 0.001, ****p < 0.0001.

### 3.10. FATP4 and HMGCL KD in primary astrocytes affect intracellular metabolism

Prompted by our *in vivo* findings, we next sought to investigate the cellular mechanisms underlying KO/KD effects on energy homeostasis. To this end, primary astrocytes were isolated from the MBH of weanling rat pups and were grown *in vitro*. siRNA-FATP4-transfected astrocytes showed an efficient and selective knockdown of FATP4 gene expression, with an approximate 70% reduction compared with controls (**Fig. 10A**). In parallel, expression of CPT1a was significantly reduced by ∼50% (**Fig. 10B**). While the latter results suggested impaired FA uptake into the mitochondria, other mitochondrial genes expression (e.g. CPT2 and CATC) were unchanged (**Fig. 10B**). While expression of cholesterol-related genes HMGCS2 and ApoE remained constant across the two treatment groups (**Fig. 10B**), siRNA-mediated FATP4 knockdown significantly reduced HMGCL mRNA levels to approximately 60% of control levels (**Fig. 10B**). Together, the findings indicate that disrupting MBH FA astrocyte transport can substantially compromise astrocytes’ ketogenic machinery.

**Figure 10.**
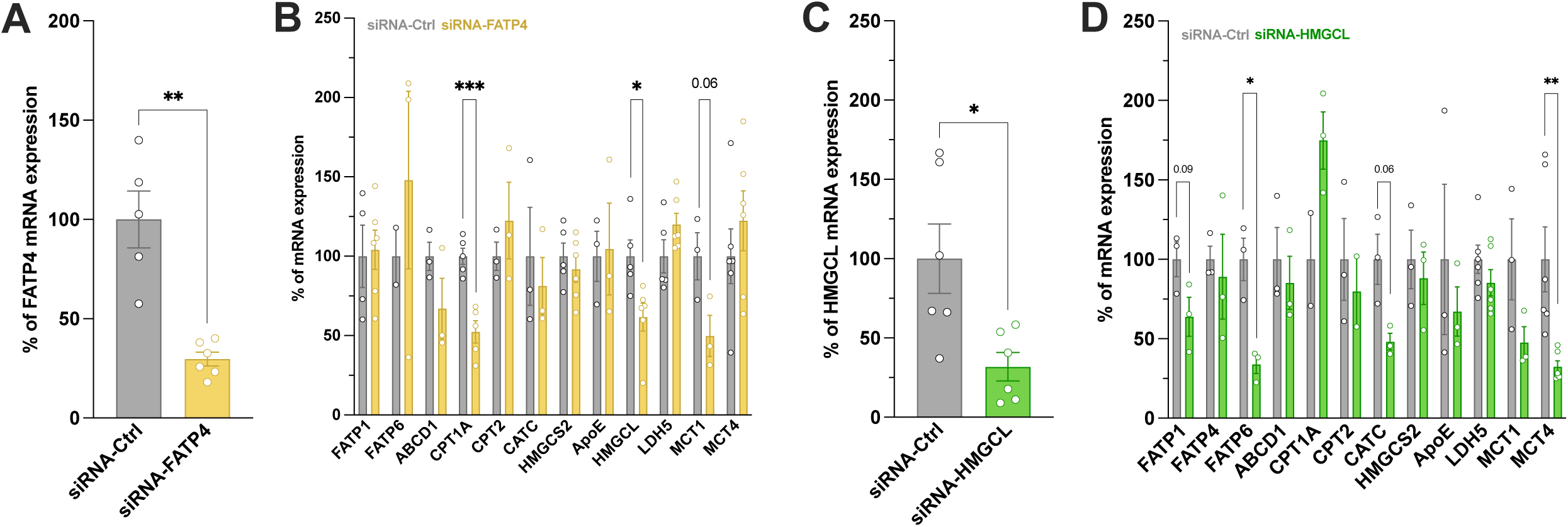
siRNA-mediated knockdown of FATP4 and HMGCL in primary MBH astrocytes induces transcriptional changes in genes involved in fatty acid metabolism. (A) FATP4 mRNA expression following transfection with siRNA-Ctrl or siRNA-FATP4. (B) Relative mRNA expression of genes involved in FA uptake and metabolism after FATP4 knockdown. (C) HMGCL mRNA expression following transfection with siRNA-Ctrl or siRNA-HMGCL. (D) Relative mRNA expression of lipid metabolism genes after HMGCL knockdown. All mRNA levels are expressed as a percentage relative to siRNA-Ctrl and were normalized to two housekeeping genes. n=3-6 per group. Data are presented as mean ± SEM and analyzed using unpaired t-tests. *p < 0.05, **p < 0.01, ***p < 0.001, ****p < 0.0001.

In addition to ketone bodies, astrocytes support neuronal energy demands through glucose metabolism and lactate production. Given that FA uptake (and potentially FA oxidation) appeared to become disrupted, we next assessed whether markers of astrocyte lactate production and shuttling were also affected. Ldh5 catalyzes the conversion of pyruvate to lactate [40], while both MCT1 and MCT4 have been implicated in the export of lactate and ketone bodies [41]. Although MCT1 showed a trend towards reduced mRNA expression (**Fig. 10B**), Ldh5 and MCT4 transcript levels were not altered by siRNA-FATP4. These data suggest that the astrocyte lactate pathway remains largely preserved in our cell culture condition despite impaired lipid metabolism.

Following the siRNA-FATP4 study, we next quantified the same gene panel in primary astrocytes treated with siRNAs targeting HMGCL mRNA (siRNA-HMGCL). It is worth pointing out that the siRNA sequences used in these experiments were identical to those employed *in vivo* for the HMGCL-KD model. The treatment reduced HMGCL mRNA levels by approximately 70% (**Fig. 10C**). Modest trends were observed in several upstream metabolic genes, particularly FATP1 (p=0.09) and CATC (p=0.06) (**Fig. 10D**). No changes were detected in MCT1 expression levels (**Fig. 10D**). In contrast, MCT4 expression was markedly downregulated by siRNA-HMGCL, a finding of particular interest that may reflect reduced export of lactate and/or ketone bodies (**Fig. 10D**).

## 4. Discussion

Accumulating evidence identifies hypothalamic astrocytes as active regulators of energy balance [9,42,43]. Building on this concept, the present study investigated the role of astrocyte lipid sensing and metabolism within the MBH in the adaptation to dietary fat excess. Using complementary loss-of-function approaches, we show that disrupting either astrocytic FA uptake through FATP4 ablation in mice or ketogenesis through HMGCL knockdown in rats exacerbates HFD-induced body weight gain. Although the phenotypes differed between models, both manipulations were associated with impaired metabolic adaptation to HFD exposure, characterized by alterations in energy expenditure and, in the HMGCL-KD model, a transient increase in caloric intake. In vitro, FATP4 depletion reduced FA-induced ketone body production and altered the expression of genes involved in mitochondrial lipid metabolism, whereas HMGCL silencing reduced MCT4 expression. Together, these findings support the concept that MBH astrocyte lipid metabolism contributes to the central mechanisms that limit the metabolic consequences of HFD exposure

Ablation of astrocytic FA transporter FATP4 in the MBH resulted in a profound disruption of energy balance during HFD feeding. FATP4-KO mice exhibited increased body weight gain and fat mass accumulation that were not driven by a sustained hyperphagic response. Although caloric intake tended to be elevated during the first days of HFD exposure, food intake subsequently remained comparable between groups. In contrast, FATP4-KO mice exhibited lower energy expenditure after chronic HFD exposure, indicating that impaired energy dissipation was likely a major contributor to their increased susceptibility to diet-induced obesity. Our data are consistent with previous studies, which demonstrate that hypothalamic astrocytes can modulate energy expenditure in the context of obesity. For instance, astrocyte-specific disruption of the circadian clock gene Bmal1 was shown to protect against diet-induced obesity by enhancing energy expenditure in response to caloric excess [44]. Conversely, chemogenetic activation of astrocytes in the paraventricular hypothalamus (PVH) of HFHS-fed obese mice suppressed whole-body energy expenditure during the dark phase [10]. Our findings extend these observations by suggesting that the ability of MBH astrocytes to take up and metabolize FAs may be required for the appropriate regulation of energy expenditure during HFD exposure.

Total energy expenditure is comprised of resting metabolic rate, locomotor activity, and adaptive thermogenesis. The reduction in energy expenditure in FATP4-KO mice occurred in the absence of changes in locomotor activity and was accompanied by a marked decrease in iBAT UCP1 protein levels, indicative of impaired thermogenic capacity [45]. The MBH regulates autonomic outflow to the BAT and serves as a critical mediator of adaptive thermogenesis, including both non-shivering thermogenesis (e.g. in response to cold) and diet-induced thermogenesis (e.g. in response to caloric excess) [46,47]. Thus, disruption of FA uptake in MBH astrocytes may be associated with the neuronal circuits underlying these thermogenic responses, thereby impairing iBAT activation during HFD feeding. Potential neuronal targets include POMC and AgRP neurons, both of which have been implicated in the regulation of iBAT sympathetic tone [48–50].

While BAT dysfunction alone may not directly cause obesity, it is thought to contribute to its progression in both rodents and humans [51]. This thermogenic defect provides a plausible explanation for the observed reduced energy expenditure. Lack of changes in locomotor activity argues against a behavioral contribution to the observed phenotype; on the other hand, core body temperature is not a reliable indicator of adaptive thermogenesis, as reductions in BAT-mediated heat production can occur without concomitant changes in whole-body temperature. Future studies incorporating direct measurements of iBAT thermogenesis or sympathetic nerve activity may be valuable in determining the extent to which adaptive thermogenesis is compromised in this specific animal model.

Beyond changes in metabolism, FATP4-KO mice also exhibited signs of glucose handling after chronic HFD exposure. Knockout mice showed elevated baseline glycemia and higher glucose excursions following oral glucose administration, resulting in an increased glucose area under the curve. Notwithstanding, the overall temporal profile of glucose clearance was similar between groups, with blood glucose concentrations returning to baseline by the end of the test, and plasma insulin levels remaining unchanged. These findings indicate that FATP4 deletion in MBH astrocytes impaired glucose tolerance without causing insulin resistance. These early disruptions in glucose regulation could, over time, increase susceptibility to type 2 diabetes and position MBH astrocyte lipid metabolism as a potential early determinant of glucose dysregulation. Astrocytes within the MBH can sense and respond to changes in glucose availability and have been implicated in the central regulation of systemic glucose homeostasis (García-Cáceres et al., 2016; Yoon and Diano, 2021). In line with this, hypothalamic astrocyte insulin signaling has been shown to regulate astrocytic coverage and synaptic organization of POMC neurons [53].

Specifically, impairments in astrocyte insulin signaling have been suggested to reduce glucose-dependent activation of POMC neurons, thereby disrupting systemic glucose homeostasis [53]. It is therefore possible that astrocyte FATP4 knockout similarly alters MBH astrocyte-POMC neuron interactions. However, because the MBH contains multiple populations of glucosensing neurons involved in the regulation of peripheral glucose levels [52], neuronal targets beyond POMC neurons (such as AgRP and NOS1 neurons) may also contribute to the hyperglycemia observed in our study. Further research will be required to establish the relative contribution of these neuronal subtypes.

The HMGCL-KD model was used to test more directly whether astrocyte ketogenesis contributes to the adaptive response to HFD feeding. In line with this hypothesis, disruption of the ketogenic HMGCL knockdown in MBH astrocytes increased HFD-induced body weight gain in rats. This phenotype partially recapitulated that of FATP4 deletion, albeit with some notable differences. In particular, the increased body weight gain in HMGCL-KD rats was not accompanied by statistically significant differences in total fat or lean mass at sacrifice. The basis for this divergence remains unclear and may reflect limitations in the sensitivity of body composition measurements or subtle changes not captured at the time of analysis. Given that individual adipose tissue depots (e.g. subcutaneous vs. visceral fat) were not quantified, we cannot rule out the possibility that depot-specific alterations may have contributed to the excess weight gain observed in HMGCL-KDs.

Unlike the FATP4-KO model, HMGCL-KD rats exhibited a transient but significant increase in caloric intake during the first three weeks of HFD exposure. This early hyperphagic response is consistent with the proposal that endogenous hypothalamic ketone body signaling contributes to the adjustment of food intake after an acute increase in dietary fat availability. In fasted rats, acute HFD consumption increases ketone body levels in the VMH before the reduction in food intake, whereas pharmacological inhibition of local ketogenesis reverses this anorectic response [15]. Similarly, local inhibition of ketone body synthesis has also been shown to promote hyperphagia in HFD-fed, diet-resistant rats that would, in steady-state conditions, reduce their food intake within three days of HFD exposure [15]. The timing of the hyperphagic response observed here coincides with the initial wave of diet-induced hypothalamic astrogliosis, which has been reported to emerge and resolve within a similar timeframe [14]. Together with acute astrogliosis, MBH astrocyte ketogenesis and ketone-dependent signaling may constitute an integral component of the adaptive response to acute HFD exposure, facilitating adjustment to increased dietary fat.

While the FATP4 knockout model displayed reduced energy expenditure in both light and dark phases, the decrease in HMGCL-KD rats was restricted to the dark (active) phase. This suggests that an early increase in caloric intake, coupled with a reduction in energy expenditure during the period of highest food consumption, may jointly be responsible for the increased body weight seen in this model. Supporting this interpretation, knockdown animals exhibited a trend towards reduced core body temperature during the dark phase at room temperature, but not under thermoneutral or cold conditions. We may thus hypothesize that MBH astrocyte ketone bodies are preferentially involved in specific components of adaptive thermogenesis, such as diet-induced thermogenesis.

The *ex vivo* iBAT analyses are consistent with this interpretation. Following cold exposure, HMGCL-KD rats showed a tendency toward reduced expression of β_3_-adrenergic receptor and UCP1, together with altered adipocyte morphology characterized by reduced cytoplasmic area and increased lipid vacuolar area [49]. These changes may reflect altered sympathetic regulation of iBAT or a reduced capacity for lipid mobilization and thermogenic remodeling. Together, these results indicate that while both knockdown and control rats respond to cold, HMGCL-KD rats may be less efficient in the regulation and fine-tuning of this response, potentially due to the combined effects of HFD and impaired MBH astrocyte ketone body production. Direct measurements of sympathetic innervation, norepinephrine turnover or mitochondrial respiration would help clarify the contribution of iBAT to the HMGCL-KD phenotype.

Several factors may explain why FATP4 deletion in mice and HMGCL knockdown in rats produced overlapping but non-identical phenotypes. First, the two manipulations target different levels of astrocyte lipid metabolism. FATP4 ablation is expected to broadly impair the uptake of long-chain FAs and thereby affect multiple downstream pathways, including lipid storage, lipid-derived signaling, mitochondrial FA oxidation, and ketogenesis [54] In contrast, HMGCL knockdown is predicted to act more selectively at the final step of ketone body production while leaving FA uptake and other aspects of lipid metabolism relatively preserved. The more sustained reduction in energy expenditure in FATP4-KO mice may therefore reflect broader impairment of astrocyte metabolic flexibility. Second, there are well-documented species-specific differences in energy balance regulation: although both food intake and energy expenditure are essential in mice and rats, rats tend to regulate energy balance through food intake, whereas mice rely more on energy expenditure adjustments [55]. Consistent with this notion, studies in the context of Roux-en-Y gastric bypass (RYGB) have shown that, in mice, body weight loss is largely driven by increases in energy expenditure rather than reductions in food intake [56,57]. These distinctions should be considered when comparing the two models, but they do not diminish the convergent observation that interference with either astrocyte FA uptake or ketogenesis within the MBH compromises metabolic adaptation to HFD.

The *in vitro* findings provide mechanistic support for a link between astrocyte FA uptake and ketogenic capacity. FATP4 silencing reduced FA-induced ketone body production in primary hypothalamic astrocytes and was associated with reduced expression of *Cpt1a and Hmgcl*. CPT1a mediates the rate-limiting transport of long-chain acyl-CoAs into mitochondria for β-oxidation [58]. Reduced *Cpt1a* expression is therefore compatible with reduced mitochondrial FA utilization, which could limit the generation of acetyl-CoA required for ketogenesis. The reduction in *Hmgcl* expression further suggests that impaired FA uptake may affect the ketogenic machinery itself. Both ketone body and cholesterol synthesis derive from acetyl-CoA and share several enzymatic steps, even though cholesterol synthesis occurs primarily in the cytosol, whereas ketogenesis is confined to the mitochondrial compartment [59,60]. As such, mitochondrial metabolic processes may be more directly affected by disruptions in FA uptake and mitochondrial substrate availability. In addition, given the essential role of cholesterol in maintaining cellular membrane integrity and function, cholesterol biosynthesis may be more tightly regulated and preserved than ketogenesis in conditions of decreased FA uptake.

HMGCL silencing in primary astrocytes resulted in a marked reduction in *Mct4* expression without substantial changes in several upstream lipid-metabolism genes. MCT4 has been implicated in the export of lactate and ketone bodies from astrocytes [41]; thus, Reduced *Mct4* expression may therefore further compromise astrocyte-to-neuron metabolic communication after HMGCL depletion, either by limiting ketone body release, lactate release, or both [61,62]. To test whether altered MCT4 expression translates into functional changes in metabolite export, lactate and ketone bodies levels could be quantified directly in the cell culture media. However, FATP4-KD could only identify a decrease in ketone production but not in lactate in response to fatty acid exposure.

Altogether, the findings of these studies identify MBH astrocytes as critical integrators of FA-derived metabolic signals within the hypothalamic circuits regulating energy balance. Strategically positioned at the interface between the circulation and nutrient-sensing neuronal networks [9,63], MBH astrocytes are equipped to sense changes in nutrient availability and initiate signaling responses necessary for the maintenance of energy homeostasis. Our complementary methodological approaches using two animal models of depletion converge on a central conclusion: disruption of astrocyte lipid metabolism within the MBH, whether through impaired FA uptake or ketone production, is sufficient to compromise adaptive responses to dietary fat, promoting the development of obesity. These results underscore the importance of intact astrocyte lipid sensing and ketone body signaling for hypothalamic control of energy balance and position astrocytes as active and essential components of the central circuitry regulating metabolic homeostasis.

## CRediT authorship contribution statement

**Keila Navarro i Batista:** Investigation, data analysis and interpretation, writing-original draft and final draft. **Patricia Kulka:** Investigation, data analysis. **Patricia Kulka:** Investigation, data analysis. **Gaia Serra:** Investigation. **Francesco Prisco:** Investigation. **Christelle Le Foll:** Conceptualization, methodology, investigation, editing, final draft, guarantor of this work and, as such, had full access to all the data in the study and take responsibility for the integrity of the data and the accuracy of the data analysis. Full datasets are available upon request. All authors approved the submitted version.

## Acknowledgments and funding

This work was funded by a Novartis grant (to C.L.F.), the Swiss National Foundation (SNF-310030_207960 and SNF-REQUIP 213511 to C.L.F.) and the Institute of Veterinary Physiology. The laboratory work was partly performed using the logistics of the Center for Clinical Studies at the Vetsuisse Faculty of the University of Zurich and the technical contributions of Dr. Petra Seebeck, Zurich Integrative Rodent Physiology. We thank Dr. Walee Chamulitrat (University of Heidelberg) for providing the FATP^fl/fl^ breeders.

