## Supplementary material for "Astrocyte fatty acid uptake and ketogenesis in the mediobasal hypothalamus coordinate adaptive metabolic responses against high-fat diet exposure in male rodents": Table S1, Table S2, Table S3, Table S4, Table S5, Table S6

**Table S1:** FATP4 genotyping primers (Microsynth AG).

| **Gene** | **Primer Sequence** |
| --- | --- |
| FATP4 | FATP4-F4 (5782312): 5’-GAA GCT ATC AGT GCT AAG CC-3’  FATP4-F10 (5782313): 5’-GAG CTT CTA TGG CAG TGA GG-3’ |

**Table S2:** Species-specific surgical parameters. Viral constructs were produced by the Viral Vector Facility of the University of Zurich.

| **Mice** | |
| --- | --- |
| Vector | **FATP4^fl: fl^:GFAP-Cre^AAV^ (v232-5; Addgene 24596):**  ssAAV-5/2-hGFAP-EGFP_iCre: WPRE-hGHp(A)  **FATP4^fl: fl^:GFAP-Control^AAV^ (v95-5; Addgene 50473):**  ssAAV-5/2-hGFAP-EGFP_WPRE-hGHp(A) |
| AAV titer during injection | 8 × 10¹¹ vg/mL |
| Anesthesia | Isoflurane (5% for induction; 1-2% for maintenance) |
| Injection volume | 300 nL/side |
| Coordinates | AP −1.4, ML ±0.3, DV −5.85 |
| Analgesia | Meloxicam (3 mg/kg; 5 mL/kg, s.c.) every 12h for 3 days  Buprenorphine (0.1 mg/kg; 5 mL/kg, s.c. ) pre-operatively |
| **Rats** | |
| Vector | **GFAP-shRNA (HMGCL)^AAV^ (vCLF4-9):**  ssAAV-9/2-hGFAP- hHBbI/ EchI [4xsh(rHmgc1)]- EGFP-WPRE-bGHp(A)  Four shRNAs against the HMGCL transcript:  GCTCTTCACCCGGAAGAATGT, GCCAAGAAGTTGTACTCAATG, GGAAATGGAATGAAGGCAAAT, GGAAATCTCTGTACGTGATTC).  **GFAP-shRNA (Control)^AAV^ (v531-9):**  ssAAV-9/2-hGFAP-hHBbI/ EchI [4x(shm/rNS)] -EGFP-WPRE-bGHp(A)) |
| AAV titer during injection | 6.9 × 10¹² vg/mL |
| Anesthesia | Ketamine (100 mg/kg; 100 mg/mL) + medetomidine (0.75 mg/kg; 1 mg/mL) + atipamezole (0.7 mg/kg; 1 mg/mL). |
| Injection volume | 500 nL/side |
| Coordinates | AP −3.2, ML ±0.33, DV −9.8 |
| Analgesia | Meloxicam (1 mg/kg; 1 mL/kg, s.c.) every 12h for 3 days  Buprenorphine (0.05 mg/kg; 1 mL/kg, s.c. ) pre-operatively |

**Table S3:** Immunostaining antibodies and dilutions.

| **Antibody** | **Host/species** | **Dilution** | **Catalogue #** |
| --- | --- | --- | --- |
| GFP | chicken | 1:500 | AB13970 (Abcam) |
| GFAP | goat | 1:1000 | AB53554 (Abcam) |
| Alexa Fluor 555 | donkey anti-goat | 1:100 | A-21432 (Invitrogen) |
| Alexa Fluor 488 | donkey anti-chicken | 1:100 | A-11039 (Invitrogen) |

**Table S4:** SYBR primers for iBAT qPCR (Microsynth AG).

| **Gene** | **Primer Sequence** |
| --- | --- |
| UCP1 | UCP1-1-F (5333188): 5’-GGG CTG ATT CCT TTT GGT CTC T-3’  UCP1-1-R (5333189): 5’.GGG TTG CAC TTC GGA AGT TGT.3’ |
| Β3-AR | b3.AR.F (5333174): 5'-ACGCTGAGGCGCAAGAGT-3'  b3-AR-R (5333175): 5'-GTAGGGCATATTGGAGGCAAAG-3' |

**Table S5:** siRNA sequences for *in vitro* FATP4 (Dharmacon) or HMGCL gene knockdown.

| **Gene** | **siRNA Sequence** |
| --- | --- |
| Slc27a4  (Dharmacon; 311839) | GACAAGGGUUUUACAGAUA  GAUUGGCCAGUGCGUAC  GAGGCUGAGUCUCGGCACA  GUACCUUCAGGAGCGGAA |
| Hmgcl  (Micronsynth) | GCUCUUCACCCGGAAGAAUGUTT  GCCAAGAAGUUGUACUCAAUGTT  GGAAAUGGAAUGAAGGCAAAUTT  GGAAAUCUCUGUACGUGAUUCTT |
| siGENOME Non-Targeting Control siRNAs (D-001210-01-05; Dharmacon) | |

**Table S6:** TaqMan assays for qPCR.

| **Pathway** | **Gene** | **TaqMan assay ID** |
| --- | --- | --- |
| Fatty Acid Transport | FATP1 (Slc27a1) | Rn00585821_m1 |
|  | FATP4 (Slc27a4) | Rn01438951_m1 |
|  | FATP6 (Slc27a6) | Custom assay: NM_001106145.1  Foward: 5’-AGTGAACCCAAAGCCATGGT-3’  Reverse 5’-GGAGGCTGGGAAGGATTTCTT-3’  FAM reporter sequence-5’-TCCAAGCAAATCTTCG-3’ |
| Lactate Shuttle | LDH5 (Ldha) | Rn00820751_g1 |
|  | MCT1 (Slc16a1) | Custom assay: NM_012716.2  Foward: 5’-CAGCTCTGACTATGATTGGCAAGTA-3’  Reverse 5’-TGCCATAGCCAGGCCATTG-3’  FAM reporter sequence-5’-CCAATGGTCGCTTCTTG-3’ |
|  | MCT4 (Slc16a3) | Rn00578115_m1 |
| Fatty Acid Metabolism | CPT1a | Rn00580702_m1 |
|  | CPT2 | Rn00563995_m1 |
|  | CATC | Rn00672611_m1 |
| Peroxisome FA Handling | ABCD1 | Rn01529709_m1 |
| Cholesterol Synthesis | HMGCS2 | Rn00597339_m1 |
|  | ApoE | Rn00593680_m1 |
| Ketogenesis | HMGCL | Rn00577753_m1 |
| Housekeeping | GAPDH | Rn01775763_g1 |
|  | RPS18 | Rn01428913_gH |
|  | Cyclophilin | Custom assay: NM_017101.1  Foward: 5’- AATGGCACTGGTGGCAAGTC -3’  Reverse 5’- GCCAGGACCTGTATGCTTCAG -3’  FAM reporter sequence-5’- TCTACGGAGAGAAATT-3’ |
